# Estimates of locus coeruleus function with functional magnetic resonance imaging are influenced by localization approaches and the use of multi-echo data

**DOI:** 10.1101/731620

**Authors:** Hamid B. Turker, Elizabeth Riley, Wen-Ming Luh, Stan J. Colcombe, Khena M. Swallow

**Affiliations:** Department of Psychology, Cornell University, Ithaca, New York, Postal Address: 211 Uris Hall, Ithaca, NY, USA; Department of Human Development, Cornell University, Ithaca, New York, Postal Address:163 Human Ecology Building, Ithaca, NY, USA; National Institute on Aging, National Institutes of Health, Baltimore, Maryland, Postal Address: 3001 S Hanover St, Baltimore, MD 21225, USA; Nathan Kline Institute for Psychiatric Research, Postal Address:140 Old Orangeburg Rd. Orangeburg, NY. 10962

**Keywords:** Locus coeruleus, multi-echo fMRI, neuromelanin-T1 imaging, resting state, intrinsic functional connectivity, norepinephrine, turbo-spin echo

## Abstract

The locus coeruleus (LC) plays a central role in regulating human cognition, arousal, and autonomic states. Efforts to characterize the LC’s function in humans using functional magnetic resonance imaging have been hampered by its small size and location near a large source of noise, the fourth ventricle. We tested whether the ability to characterize LC function is improved by employing neuromelanin-T1 weighted images (nmT1) for LC localization and multi-echo functional magnetic resonance imaging (ME-fMRI) for estimating intrinsic functional connectivity (iFC). Analyses indicated that, relative to a probabilistic atlas, utilizing nmT1 images to individually localize the LC increases the specificity of seed time series and clusters in the iFC maps. When combined with independent components analysis (ME-ICA), ME-fMRI data provided significant gains in the temporal signal to noise ratio relative to denoised single-echo (1E) data. The effects of acquiring nmT1 images and ME-fMRI data did not appear to only reflect increases in power: iFC maps for each approach only moderately overlapped. This is consistent with findings that ME-fMRI offers substantial advantages over 1E data acquisition and denoising. It also suggests that individually identifying LC with nmT1 scans is likely to reduce the influence of other nearby brainstem regions on estimates of LC function.

**Highlights:**

- Manual tracing of locus coeruleus increased specificity of seed time series
- Manual tracing of locus coeruleus increased specificity of intrinsic connectivity
- Multi-echo fMRI increased temporal signal-to-noise ratio compared to single-echo fMRI
- Connectivity maps across methodologies overlapped only moderately
- Measurement of LC function benefits from multi-echo fMRI and tracing ROIs

## Introduction

The locus coeruleus (LC) is a pair of small, cylindrical nuclei located in the brainstem near the fourth ventricle (4thV). As the main site for the synthesis of the neuromodulator norepinephrine (NE), the LC influences many aspects of cognition and autonomic regulation (e.g., Aston-Jones, Gonzalez, & Doran, 2007). Because of its small size and location, however, investigations of LC function in humans using functional magnetic resonance imaging (fMRI) face a unique set of challenges for confidently localizing and isolating signals associated with neural activity in the LC. This paper investigates the effectiveness of two data acquisition and processing approaches in mitigating those problems: neuromelanin-weighted T1 (nmT1) imaging to localize the LC in each individual (Sasaki et al., 2006) as well as multi-echo functional magnetic resonance imaging (ME-fMRI) to increase blood oxygen level dependent (BOLD) contrast and reduce non-BOLD artifacts (Kundu, Inati, Evans, Luh, & Bandettini, 2012). To do so, we characterize differences in estimates of the intrinsic functional connectivity (iFC) of the LC when these approaches are and are not used.

Accurately characterizing LC function is important because it has implications for our understanding of a wide range of cognitive processes and neuropsychological disorders. Despite its small size, the LC projects to most of the central nervous system, excluding the striatum, and is the primary source of NE in the brain (Berridge & Waterhouse, 2003; Jones & Moore, 1977; Jones & Yang, 1985; Samuels & Szabadi, 2008). The LC contains sub-populations of cells that project to different regions of the brain, such as the brainstem, spinal cord, medial prefrontal cortex, hippocampus, septum, and amygdala (Chandler, Gao, & Waterhouse, 2014; Schwarz & Luo, 2015), and has particularly dense projections to sensory and motor regions (Loughlin, Foote, & Bloom, 1986; Schwarz & Luo, 2015). LC projections also show large amounts of branching along the anterior-posterior axis in neocortex that may be driven by the shared function of target regions (Loughlin, Foote, & Fallon, 1982; Aston-Jones & Waterhouse, 2016). In turn, the LC receives input from the brainstem, hypothalamus, the central nucleus of the amygdala, the anterior cingulate cortex, and the orbitofrontal cortex (Aston-Jones et al., 1991; Luppi, Aston-Jones, Akaoka, Chouvet, & Jouvet, 1995). This circuitry allows the LC to regulate autonomic states and task engagement to influence how neural systems respond to behaviorally relevant events (Aston-Jones, Rajkowski, & Cohen, 1999; Glennon et al., 2019; Greene, Bellgrove, Gill, & Robertson, 2009; Nieuwenhuis, Van Nieuwpoort, Veltman, & Drent, 2007; Mohanty, Gitelman, Small, & Mesulam, 2008). Although the precise mechanisms by which LC activity accomplishes this task are still being investigated (Mather, Clewett, Sakaki, & Harley, 2016; Uematsu, Tan, & Johansen, 2015; Schwarz & Luo, 2015), NE changes the signal-to-noise ratio in brain regions involved in perceptual processing (Foote, Freedman, & Oliver, 1975; Berridge & Waterhouse, 2003), promotes memory consolidation and retrieval (Sara, 2009; Grella et al., 2019; Swallow, Jiang, & Riley, 2019), and may influence the integration of networks across brain regions (Shine et al., 2016). The importance of the LC for basic cognitive processing has been further reinforced by findings that there is an estimated 40% decline in cell number in the LC over the lifespan (Vijayashankar & Brody, 1979) which is accompanied by a decline in cognitive performance (Mather & Harley, 2016).

### Localizing human LC and measuring its neuromelanin content using MRI

Given the importance of the LC for cognitive and neural processing across the lifespan, a growing number of studies have begun to investigate its structure and function using magnetic resonance imaging (MRI). However, the size and location of the LC nuclei pose unique challenges to identifying the LC in these scans. LC nuclei are located in the upper portion of the pons, have between 22,000 and 51,000 neurons depending on one’s age (Mann, Yates, & Hawkes, 1983; Mather & Harley, 2016), and range in size between 31,000 and 60,000 µm^3^ (Mouton, Pakkenberg, Gundersen, & Price, 1994) with a within-plane diameter of only 2.5 mm (Fernandes, Regala, Correia, Gonçalves-Ferreira, 2012). In standard T1-weighted images, the LC and the surrounding regions appear relatively homogenous, making identification of voxels that contain the LC difficult. This is exacerbated by its small size and variability in its location (Keren, Lozar, Harris, Morgan, & Eckert, 2009). To help address this problem, neuroimaging sequences have been developed that increase the contrast for regions containing neuromelanin (nmT1 imaging), a pigment found in the LC (Sasaki et al., 2006). Neuromelanin is an iron containing product of catecholamine synthesis that causes paramagnetic T1 shortening (Enochs, Petherick, Bogdanova, Morh, & Weissleider, 1997; Sulzer, et al., 2018; Wakamatsu, Tabuchi, Ojika, Zucca, Zecca, & Ito, 2015). These paramagnetic effects can be leveraged by nmT1 imaging to increase the contrast of neuromelanin containing voxels to allow for the visualization and localization of the LC in structural MRI scans (Keren et al., 2009). Because regions that contain more neuromelanin appear brighter in nmT1 images, it also provides an *in vivo* measure of the amount of neuromelanin in the LC (and the other neuromelanin containing region in the brainstem, the substantia nigra) (e.g., Keren et al., 2009; Keren et al., 2015; Liu et al., 2019).

However, the majority of studies on LC function do not make use of nmT1 imaging to localize the LC. Instead, they rely either on previously published coordinates, probabilistic atlases (such as by Keren et al., 2009) or use exploratory whole-brain analyses (for a review, see Liu et al., 2017). But because the precise location and shape of a brain region can vary across individuals, defining regions of interest (ROIs) at the group level risks missing the region in the individual and capturing signal from other areas. This can result in group level ROIs producing less reliable findings than individually defined ROIs (e.g., Swallow, Braver, Snyder, Speer, & Zacks, 2003). Therefore, coordinates from other studies or probabilistic LC ROIs, while likely to capture the LC, may also capture other nearby brainstem structures such as the inferior colliculus, the nucleus incertus, or other parts of the ascending reticular activating system that also play a role in arousal, orienting, and learning (e.g., Ryan, Ma, Olucha-Bordonau, & Gundlach, 2011). Accordingly, one of the main interests in the current study was to compare the effects of using nmT1 imaging versus using a probabilistic atlas on one’s eventual characterization of LC function.

Concerns about an ROI capturing activity of nearby anatomical regions or cerebrospinal fluid (CSF) signal are further exacerbated by a commonly employed data preprocessing step: spatial blurring. Blurring reduces the effects of noise by averaging over nearby voxels. It also increases the likelihood that the ROI includes the signal of interest (Mikl et al., 2008). However, for small regions like the LC, blurring with a large kernel may inadvertently reduce one’s ability to detect its activity. This is particularly problematic for regions, like the LC, that are close to large sources of physiological noise or changes in tissue. In these cases, blurring may introduce additional physiological noise into the signal unless measures are taken to preclude this possibility (e.g., by removing voxels from the fourth ventricle prior to blurring). When it comes to investigations of the LC, data blurring has varied widely across studies (Liu et al., 2017) and there has been no systematic investigation of whether blurring, even by modest amounts, alters estimates of LC function and connectivity.

### Estimating LC activity: multi-echo versus single-echo functional MRI

Previous investigations of LC function have used single-echo functional magnetic resonance imaging (1E-fMRI), which measures the MR signal in each voxel once per volume acquisition. By measuring the MRI signal two or more times per acquisition, multi-echo (ME) fMRI offers two potential advantages over 1E-fMRI that could be particularly effective at reducing the impacts of non-BOLD noise and signal drop-out on estimates of LC function: maximization of BOLD contrast through the optimal combination of echoes during data analysis and ICA denoising.

With 1E-fMRI, the time at which the signal is measured (TE) is selected to maximize BOLD contrast for the brain as a whole, balancing variability in the T2* signal decay rates across the brain that arise from regional differences in tissue composition and signal drop-out (e.g., Cho & Ro, 1992; Park, Ro, & Cho, 1988). Because ME-fMRI measures the MR signal at multiple points during each acquisition, it can be used to estimate initial signal intensity and the rate at which the signal decays (T2*) at every voxel in the volume. This information can be used to create volumes that optimally weigh and combine the TEs for each individual voxel. Rather than choosing one TE for the entire brain, ME-fMRI can be used to effectively estimate the signal at the optimal TE for a given voxel and brain region after the data have been acquired (Kundu, Voon, Balchandani, Lombardo, Poser, & Bandettini, 2017). This approach has been used to improve BOLD contrast in regions of the brain that are subject to signal-dropout in typical 1E-fMRI, such as the ventromedial prefrontal cortex and nucleus basalis, without sacrificing contrast in other brain regions (Markello, Spreng, Luh, Anderson, & De Rosa, 2018).

Estimation of initial signal intensity and T2* decay also affords ME-fMRI data with a theoretically motivated approach to denoising BOLD data. Changes in BOLD activity are dependent on the TE, while artifactual effects on MR signal are TE-independent (Kundu et al., 2013). ME independent components analysis (ME-ICA) takes advantage of these differences in order to identify and remove the contributions of non-BOLD signal components, such as head motion and physiological noise, from the data. This approach has been shown to significantly reduce the effects of head motion and other noise sources on BOLD data (Power et al., 2018). In contrast, denoising of 1E-fMRI data often requires removing observations that are likely to be contaminated by motion (e.g., by scrubbing) or by estimating and regressing out noise from motion and physiology (see, e.g., Dipasquale et al., 2017). For example, with RETROICOR, variance associated with cardiac phase and respiration is removed from the data using regression (Glover, Li, & Ress, 2000). Other approaches (e.g., ANATICOR; Jo, Saad, Simmons, Milbury, & Cox, 2010) estimate non-neural contributions to the BOLD signal by regressing out signal captured in nearby white matter voxels or CSF. Principle components and independent components analysis (e.g., ICA-AROMA; Pruim, Mennes, Van Rooij, Llera, Buitelaar, & Beckmann, 2015) can also be used to remove sources of noise from 1E-fMRI data. These methods are often employed in conjunction with the inclusion of first- and second-order motion regressors in the statistical analyses. However, when directly compared to ME-ICA, these methods have been shown to be less effective at reducing noise (reflected in greater temporal signal to noise ratio and lower DVARS; Lombardo et al., 2016; Markello et al., 2018; Dipasquale et al., 2017).

Previous studies investigating the iFC of the LC have not consistently accounted for cardiac and respiratory effects, and few have reported correcting for motion (Liu et al., 2017). For example, Murphy, O’Connell, O’Sullivan, Robertson, & Balsters (2014) explicitly report applying RETROICOR. Krebs, Park, Bombeke, & Boehler (2018) instead do not denoise their data from physiological noise but compare activity in the LC to neighboring regions and conclude from the dissimilarities that their LC activity is not primarily driven by noise. Because physiological and motion-related noise can cause spurious correlations in iFC maps (Power, Barnes, Snyder, Schlagger, & Petersen, 2012), steps to reduce their impact on the data are needed to gain a better understanding of LC function. This is particularly important for the LC because of its size and location near the 4thV, and increased likelihood that it will contain signals from CSF. In fact, previous neuroimaging studies showing task-related activity in the vicinity of the LC have been called into question precisely because of this problem (Astafiev, Snyder, Shulman, & Corbetta, 2010).

A previous investigation of the use of ME-ICA with a pair of small brain regions in the human basal forebrain, the nucleus basalis of Meynert and the medial septum (Markello et al., 2018), suggest that ME-ICA may also provide significant advantages over 1E-fMRI approaches when investigating the iFC of the LC. Like the nucleus basalis of Meynert and the medial septum, the small size and location of the LC near the boundary between tissue and CSF make it particularly important to account for signal dropout from local field inhomogeneities and to remove noise from non-neural sources. Therefore, to investigate whether ME-fMRI offers a benefit above and beyond traditional single-echo scans, this study examines differences in the estimated functional characteristics of the LC when using ME-fMRI relative to 1E-fMRI.

### The current study

Human neuroimaging of the LC has been hampered by several related issues: localizing the LC, isolating LC activity from other nearby regions, and removing the effects of noise from motion and physiology. In this study, we investigate whether using nmT1 images, ME-fMRI, and conservative amounts of data blurring offer appreciable advantages to efforts to characterize LC function. We also briefly describe a method for standardizing nmT1 signal intensity using the whole brainstem, as previously reported approaches are inconsistent in their use of a reference region. Intrinsic functional connectivity analyses of resting state data formed the basis for comparing these different approaches (Biswal, Yetkin, Haughton, & Hyde, 1995; Van den Heuvel & Pol, 2010). Although previous studies have characterized iFC of the LC using resting state data (e.g. Murphy et al., 2014; Song, Kucyl, Napadow, Brown, Loggia, & Akeju, 2017; Zhang, Hu, Chao, & Li, 2016), there is little published data investigating the impact of different methodological and analytical approaches on estimates of LC function (but for a review of approaches used, see Liu, et al., 2017). The current paper meets this need and suggests that different localization, data acquisition, and pre-processing approaches can in fact lead to different characterizations of LC function and connectivity.

## Methods

### Participants

A total of 20 right-handed participants (14 female, 6 male, 19-40 years old, M = 21.05, SD = 4.57) completed the task. Participants were screened for non-MRI compatible medical devices or body modifications (e.g., piercings, implants), claustrophobia, movement disorders, pregnancy, mental illness, use of medication affecting cognition, and color blindness. All participants provided consent at the start of the session, were debriefed at the end, and all procedures were approved by the Cornell University Institutional Review Board.

### MRI Acquisition

Magnetic resonance imaging was performed with a 3T GE Discovery MR750 MRI scanner and a 32-channel head coil at the Cornell Magnetic Resonance Imaging Facility in Ithaca, NY. Participants laid supine on the scanner bed with their head supported and immobilized. Ear plugs, headphones, and a microphone were used to reduce scanner noise, allow the participant to communicate with the experimenters, and to present auditory stimuli during the tasks. Visual stimuli were presented with a 32” Nordic Neuro Lab liquid crystal display (1920×1080 pixels, 60 Hz, 6.5 ms g to g) located at the back of the scanner bore and viewed through a mirror attached to the head coil. Pulse oximetry and respiration were recorded throughout all scans.

Anatomical data were acquired with a T1-weighted MPRAGE sequence (TR = 7.7 ms; TE = 3.42 ms; 7° flip angle; 1.0 mm isotropic voxels, 176 slices). A second anatomical scan utilized a neuromelanin sensitive T1-weighted partial volume turbo spin echo (nmT1) sequence (TR = 700 ms; TE = 13 ms; 120° flip angle; 0.43 x 0.43 mm in-plane voxels, 10 interleaved 3.0 mm thick axial slices; adapted from Keren, et al., 2009). Slices for the nmT1 volume were oriented perpendicular to the brain stem to provide high resolution data in the axial plane, where the dimensions of the LC are smallest, and positioned to cover the most rostral portion of the pons.

Participants completed one resting state scan with eyes open and the lights on during multi-echo echo planar imaging (612 s; TR=3.0 s; TEs=13, 30, 47 ms; 83° flip angle; 3.0 mm isotropic voxels; 46 interleaved slices). The display was set to a uniform light grey background throughout the scan. Participants were told to keep their eyes open and remain awake throughout the scan, but were free to move their eyes and blink as needed. Eye movements, blinks, and pupil size were recorded with an Eyelink 1000 Plus MRI Compatible eye tracker (SR-Research, Ottawa, Ontario, Canada). However, because participants were free to blink and move their eyes as needed, the recordings were too noisy to analyze with confidence.

### Region of Interest Identification

#### Anatomically Defined Regions

Anatomical data were submitted to FreeSurfer’s segmentation and surface-based reconstruction software (recon-all; v 5.3; http://surfer.nmr.mgh.harvard.edu/; Dale, Fischl, & Sereno, 1999; Fischl, Sereno, & Dale, 1999) to label each individual’s anatomy. Labels for cortical gray matter (GM), ventral medial prefrontal cortex (vmPFC), fourth ventricle (4thV), hippocampus (HPC), primary visual cortex (V1), brainstem, precentral gyrus (motor cortex, MC), and transverse temporal gyrus (auditory cortex, AC) were extracted and converted to volumetric ROIs using FreeSurfer and AFNI (Cox, 1996) tools. A white matter (WM) ROI was similarly created, but eroded using AFNI’s 3dmask_tool (dilate_input −2) to ensure that the ROI did not extend into nearby regions.

#### Locus Coeruleus

Voxels that were likely to include the locus coeruleus were identified in two ways: using the probabilistic Keren atlas, and the nmT1 images. ROIs that were based on the Keren atlas were defined using the binary 1 SD and 2 SD masks provided by those authors (referred to here as K1 and K2 ROIs, respectively; Keren et al., 2009) in the MNI atlas (T1 MNI-152 0.5 mm iso-voxel). To provide the most conservative comparison between the Keren atlas and the nm-based LC ROI, analyses focused primarily on the K1 ROI. The Keren atlas is based on nmT1 images (3 x 0.4 x 0.4 mm axial slices) of 44, right-handed, healthy adults, aged 19-79. The final probabilistic atlas was based on the group means and standard deviations of the highest intensity voxels on the left and right in each axial slice (for more details see Keren, et al., 2009). Individual, hand-traced LC ROIs (referred to as the LC ROI from here on) for each participant in the present experiment were defined using the nmT1 image, with the same scanning parameters as used for the Keren atlas (Figure 1). Prior to tracing the ROIs, the nmT1 image was processed by normalizing intensity within each slice. Then, brainstem and 4thV voxels were extracted from the nmT1 images. Masks for these regions were created by aligning the individual’s anatomical parcellation from Freesurfer to the nmT1 image using a 12 parameter affine transformation. These parameters were calculated by aligning the participant’s MPRAGE volume (with skull, for additional landmarks) to the nmT1 volume (AFNI’s align_epi_anat.py; Cox & Jesmanowicz, 1999; Saad, Glen, Beauchamp, Desai, & Cox, 2009). To normalize the image, the mean signal intensity of voxels within the brainstem mask was subtracted from each masked slice of the nmT1 image.

**Figure 1.**
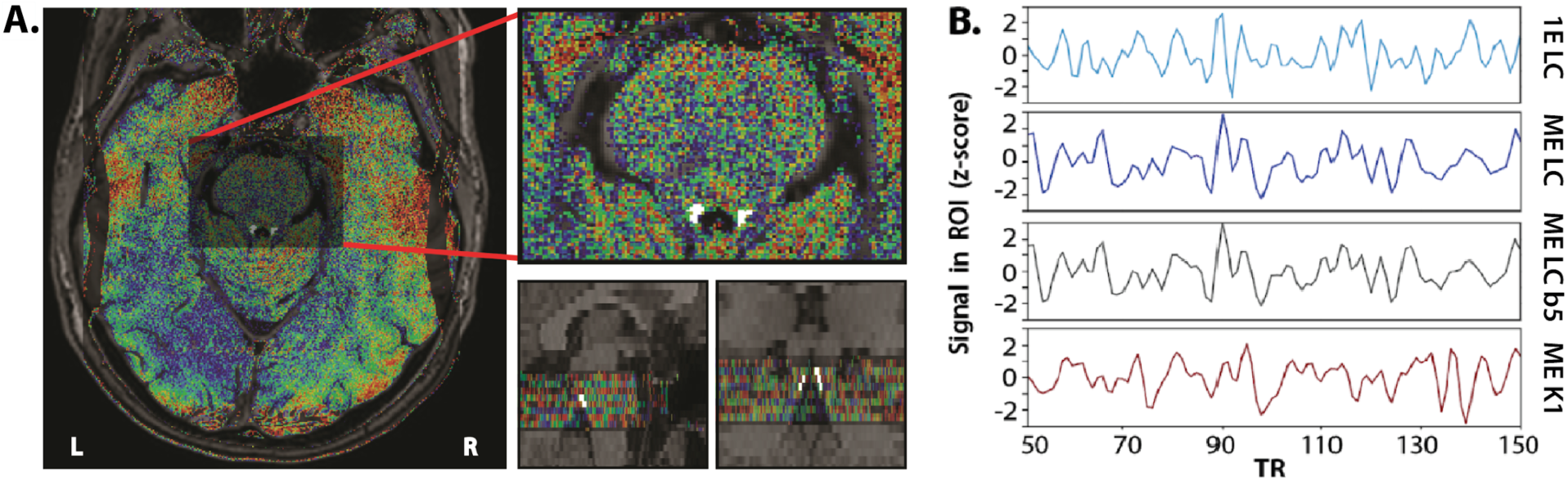
Visualization of the ROI creation and seed extraction process. **A.** Individual MPRAGE scans (including skull) were aligned to the normalized nmT1 image. Values in the nmT1 image are shown in false color (FSL’s jet palette: low values in blue, high values in red, range = 10-80) overlaid on the aligned MPRAGE showing the whole brain in an axial plane (left image), zoomed in on the pons (upper right), and in sagittal (lower middle) and coronal (lower right) planes. Two researchers independently traced the LC according to the protocol described in the methods. The overlap of their ROIs (shown in white) was resampled to 3mm voxels and aligned with the native MPRAGE. **B.** Excerpts from one individual’s seed time series (from acquisition numbers 50 to 150; z-scored) of the ME LC without blurring, 1E LC without blurring, ME LC with blurring up to 5mm FWHM of the seed (ME LC b5), and ME K1 seed without blurring. LC seeds were extracted after the ROI was aligned with the native MPRAGE. K1 seeds were extracted after warping to the MNI atlas.

The LC was manually defined in FreeView (https://surfer.nmr.mgh.harvard.edu) on the corrected and masked nmT1 images by two independent raters (authors HBT and ER) using the following criteria (adapted from Tona et al., 2017):

1. The corrected nmT1 image was viewed in false color and overlaid on the aligned MPRAGE image. Brush size was 1.
2. Visual contrast was equated across participants using a predefined range for the false color palette (Jet - minimum: 10, maximum: 80). Pixel intensity threshold for inclusion in the LC (only green, yellow, orange and red pixels could be included) was agreed upon prior to drawing the ROIs.
3. Two landmarks were used to locate the most rostral portion of the LC: the lower boundary of the inferior colliculus (seen in sagittal view) and the upper boundary of the 4thV (seen in axial view). The rostral end of the LC was in the first axial slice caudal to the inferior colliculus in which two distinct hyperintensities on the edge of the 4thV were visible in the nmT1 image.
4. The LC was defined starting on this slice and moving caudally through the pons. Hyperintensities in more caudal slices were included in the LC only if they were both sufficiently bright and connected to hyperintensities in the next most rostral slice, verified in sagittal and coronal views.
5. Tracing was informed by the assumption that each LC nucleus would appear as a roughly circular shape within each axial slice, resulting in bilateral, roughly cylindrical regions. In addition, the LC was assumed to be solid and contain no holes. If a voxel was surrounded in-plane by voxels included in the LC, then that voxel was also included.
6. In some cases, a second, more medial hyperintensity was found near the rostral part of the LC. This region was taken to be part of the trochlear nerve and was not included in the LC ROI. Avoiding this secondary hyperintensity prevented the LC ROI from having a very oblong shape or extending to the midline at any point.

An individual’s final LC ROI consisted of voxels included in both raters’ ROIs, after removing any voxels that overlapped with 4thV. 4thV was defined starting from the most rostral axial slice in which the cerebral aqueduct began to widen, until the most caudal axial slice in which the ventricle appeared dark and had an inverted-U shape.

Seeds for the iFC analyses of the LC ROIs (drawn on the nmT1; Figure 1) were acquired by aligning the LC ROIs to the functional data that were aligned to the native MPRAGE. A probabilistic map illustrating the location and distribution of the traced LC in all participants was created by registering each person’s LC ROI to MNI space (MNIa_caez_N27). Alignment parameters were calculated for each individual by combining the inverse of the 12 parameter affine transformation that aligned the native MPRAGE and nmT1 images, the 12 parameter affine transform from the individual MPRAGE to the MNI standard, and a nonlinear transform from the individual’s native MPRAGE to MNI standard calculated by 3dQwarp. The alignment parameters were then simultaneously applied to the LC ROI using 3dNwarpApply. Successful alignment of the brainstem in the individual MPRAGE and the MNI template was visually confirmed. Similar to previous findings (Tona et al., 2017), linear transformations were inadequate for registering a small brainstem ROI to a standardized space: with affine registration only, many LC ROIs overlapped the 4thV or other parts of the pons. Aligned LC ROIs were thresholded to remove non-zero voxels introduced by the nonlinear warping process. Finally, each voxel value was divided by the number of participants to create the probabilistic map (Figure 2).

**Figure 2.**
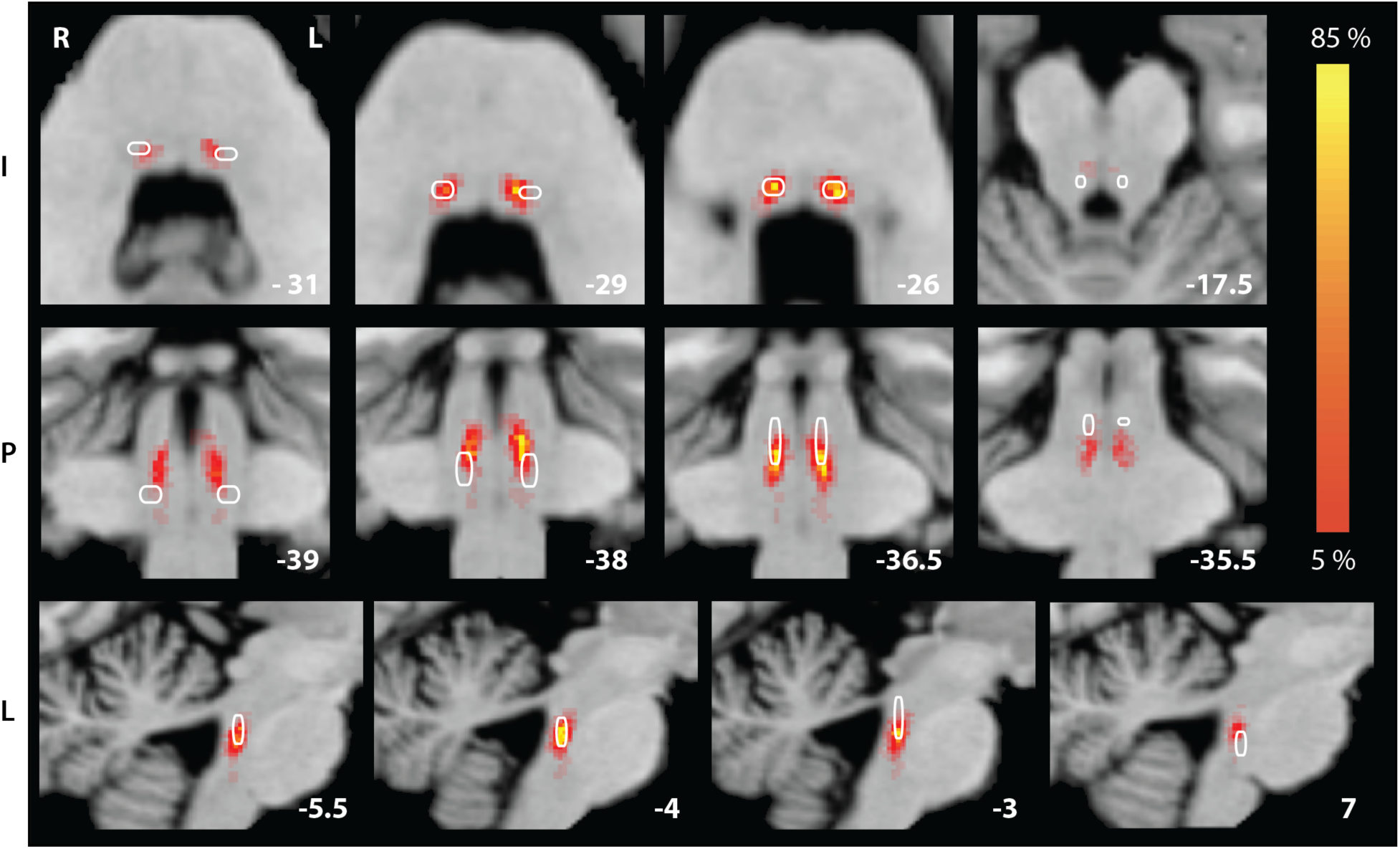
Hand-traced LC atlas (heat; percentage of participants in the current study) and binary K1 atlas (white outline) in MNI. Slices were selected to best visualize similarities and differences between the LC heat map and the K1 ROI. In analyses, individual traced LC ROIs were applied to data aligned to the individual’s native anatomy, but warped here to MNI for aggregation and comparison to the Keren atlas. Coordinates in LPI.

#### Pontine Tegmentum

Within axial slices of the participant’s MPRAGE (as aligned to the nmT1), the pontine tegmentum (PT) ROI was defined as a circular ROI 10 voxels in diameter centered at the midline of the brainstem, ventral to the fourth ventricle, and approximately equidistant from the left and right LC ROIs, to define the vertex of an equilateral triangle. The PT ROI was defined only in slices that also contained an LC ROI (see also Clewett, Lee, Greening, Ponzio, Margalit, & Mather, 2016).

### LC Neuromelanin Intensity

To compare neuromelanin intensity across individuals, rescaling parameters were obtained from the corrected nmT1 image: the mean, standard deviation, a minimum value (*min = M-3*SD*), and a maximum value (*max = M+3*SD*) were calculated using non-LC voxels from axial slices in which LC was defined. Using these non-LC voxels, all voxels in the corrected nmT1 image were then rescaled using the equation 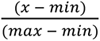, resulting in an image with a unit scale equal to 6 standard deviations of the non-LC voxel signal intensities. Because the LC was defined as hyperintensities that were excluded from the scaling parameters, values greater than 1 were expected within the LC. The rescaled intensity values within the LC ROIs were then averaged to obtain the mean intensity for each participant.

### MRI Data Pre-Processing

To evaluate the effects of different denoising procedures on data quality and the resulting iFC maps, four single-echo pre-processing pipelines and one multi-echo pre-processing pipeline were used. Pre-processing of ME resting state data were modeled after the procedures outlined in Jo et al. (2013) and Markello et al. (2018). The 1E and ME pipelines were matched as closely as possible. We first describe the pipeline used for ME-ICA denoising, followed by the variations 1E pipeline, blurring, and nuisance regressor extraction).

For ME data sets with *ME-ICA-denoising*, the standard ME-ICA pipeline (meica.py, Version 2.5, beta 9; tedana.py, Version 2.5 beta 9; t2smap.py, Version 2.5 beta 6; Kundu et al., 2012; Kundu et al., 2013) was implemented using the following steps. First, the MPRAGE volume was skull stripped using FSL BET (b = 0.25). Second, the obliquity of the anatomical volume was matched to the EPI time series. Third, motion was estimated using the first echo time series using 3dvolreg with the third volume as the target. Fourth, all EPI data were despiked and slice time acquisition differences were corrected using 3dTshift. Fifth, for each echo time series, the first two volumes were dropped and the remaining EPI data were registered to the third volume. Sixth, the three echoes were concatenated and processed by t2smap.py to generate the baseline intensity volume (s0), the T2* relaxation map, and the optimal combination volume time series (OCV). Seventh, registration and alignment transforms were applied to the EPI data in one step to align the data with the individual anatomical volume in its original acquisition space. Eighth, EPI data were denoised using ME-ICA to identify and separate BOLD components from non-BOLD components (Kundu, et al., 2013). The BOLD components were recombined to create the denoised data sets used in the analyses. Ninth, EPI data were warped to MNI space to be used with the K1 and K2 atlas; a copy of the EPI aligned to the MPRAGE was kept to be used with the individually defined LC ROIs. Because global signal regression was not used, nuisance regressors capturing unaccounted for physiological and motion related noise (Power et al., 2018) (demeaned motion) were then obtained by averaging the time series of each voxel within the WM and 4thV ROIs. All regressors were obtained on unblurred data, to avoid contaminating the regressor with signal from neighboring areas (Jo et al., 2013). Spatial blurring to 5 mm FWHM was then performed (3dBlurtoFWHM). Finally, the data were bandpassed (0.01 < *f* < 0.1) and nuisance regressors were removed, all in one step, with 3dTproject (cenmode NTRP).

1E data were denoised in four different ways. In each case, only the second echo of data was analyzed. For the *WM-regression-only denoising* 1E dataset, pre-processing skipped steps 6 and 8, after which only WM signal was included as a nuisance regressor alongside the standard removal of demeaned motion and their first derivatives. The *RICOR+WM-denoising* pipeline was the same as the WM-regression-only pipeline with the addition of RETROICOR denoising: respiration and pulse oximetry data were used to generate per slice regressors using AFNI’s retroTS.py (Glover et al., 2000). RETROICOR (shortened to RICOR) statistically removes physiological noise from the EPI data between steps 3 and 4 using 3dREMLfit. An additional pipeline examined the effectiveness of removing signal in the 4thV: the *4thV+WM-denoising 1E* data was similar to the WM-regression-only pipeline, with the addition of the 4thV time series as a nuisance regressor. The fourth and final 1E-denoising pipeline (RICOR+4thV+WM-denoising) included all of the aforementioned denoising procedures.

To examine the effects of blurring prior to seed extraction we extracted the time series within the LC, K1, K2 and PT ROIs both before (native blur) and after (b5) the data were blurred to 5 mm FWHM.

### Intrinsic Functional Connectivity Analyses

For first level functional connectivity analyses, seeds were created by averaging the time series of voxels within the ROIs. LC seeds were extracted after aligning the data to the individual MPRAGE, while K1 and K2 seeds were extracted after aligning the data to the MNI template. Seeds were then correlated with the timeseries of each voxel in the blurred data set in MNI space. The Fisher r-to-z transform was then applied to the correlation maps to produce z-maps for the second level analysis. In the second level analysis, voxel-wise t-tests (3dTtest++) were performed to test each pipeline against zero and the statistical maps were controlled for multiple comparisons using the false discovery rate (FDR) (Genovese, Lazar, & Nichols, 2002). First and second level analyses were performed for each of the denoised data sets to produce functional connectivity maps that varied along several factors: 1E vs. ME (*data type*), LC vs K1 seeds (*ROI*), and no-blur vs. blur to 5 mm (b5) prior to seed extraction (*blur*).

### Data and Code Availability Statement

Because participants did not consent to their data being made public, the data from this study are not publicly available. However, the corresponding author can provide them upon request. Analyses used publicly available software (AFNI, FSL, meica.py). Annotated analysis scripts for AFNI and R can be found on the GitHub of author HBT (https://github.com/HamidTurker/estimatesoflcfunction).

## Results

### Summary of Comparisons and Analyses

The aforementioned procedures allowed us to examine the effects of several data acquisition or pre-processing choices on data quality and functional connectivity maps of putative LC regions. Analyses were carried out in two phases. The first phase focused on assessing changes to the temporal signal to noise ratio (tSNR) following ME-ICA on ME-fMRI data as well as the various forms of 1E data denoising. To preview the results, the initial analyses suggested that RICOR combined with WM and 4thV regression offer appreciable increases in the tSNR of the 1E data. Therefore, in the second phase, where we characterize the intrinsic functional connectivity of the LC, only the RICOR+4thV+WM-denoised 1E data and ME-ICA denoised data are used. Both datasets included WM and 4thV nuisance regressors.

### Locus coeruleus localization and intensity measurement

To characterize the location and reliability of the LC ROIs defined on the nmT1 images (shown in white in Figure 1A), we measured inter-rater agreement and inter-individual differences in LC size and location. The average Dice similarity coefficient between the raters’ ROIs was 0.81 (range 0.44 - 1.00). In MNI space, the average location of the center of mass of the right and left LC were (LPI: −5.26, 37.15, −25.91) and (LPI: 3.67, 37.30, −26.36), respectively. The LC had an average total volume of 37.5 ± 11.8 mm^3^, with an average length of 9.28 ± 2.28 mm. The volume of the right ROI was significantly larger than the left ROI (right volume 21.90 ± 7.14 mm^3^, left volume 15.56 ± 5.43 mm^3^, paired t-test, t(20) = 6.35, *p* < 0.001). The length did not differ significantly between the right and left ROIs. These values show good agreement with those of postmortem studies on “core” LC volume (German, Walker, Manaye, Smith, Woodward, & North,1998, Fernandes et al., 2012). In addition, the group-level heat map visually overlapped with the K1 ROI (Figure 1), though the average Dice coefficient between the individual LC ROIs and the K1 ROI was low (M = .195, SD = .070, range = 0 - .311). Accordingly, the K1 ROI consisted of different, and more, voxels than the traced LC ROIs for each individual.

LC intensity was standardized across participants using the mean and standard deviation of non-LC voxels within the brainstem. This ensured that the relative brightness of LC voxels was measured regardless of the overall image intensity, or the intensity of a particular anatomical region (e.g., the pontine tegmentum, which also shows an effect of aging, Clewett et al., 2016; Keren et al., 2009). The average intensity, in units of standard deviation, within the right and left LCs respectively were found to be 0.81 (SD = 0.05) and 0.80 (SD = 0.06). The difference between them approached significance, paired t-test, t(20) = 1.91, *p* = 0.071.

### Quality assessment of ME-ICA denoising procedure

To assess the effectiveness of the different denoising procedures, we calculated the tSNR using AFNI’s 3dTstat (cvarinvNOD). tSNR was calculated for the GM, HPC, V1, vmPFC, MC, AC, and LC ROIs (Table 1).

**Table 1.**
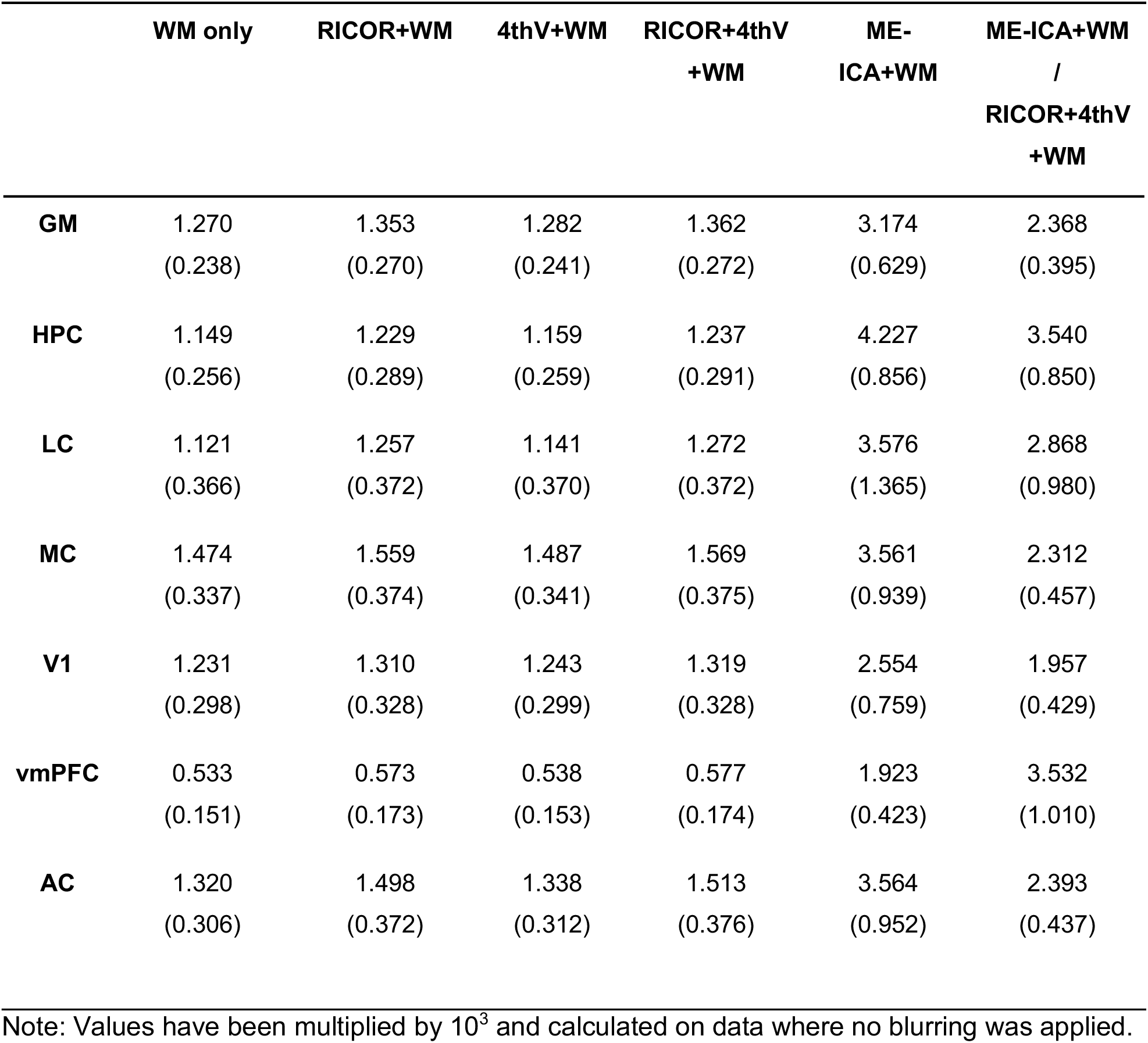
Means and standard deviations (in parentheses) of tSNR in each region of interest following the five denoising procedures.

|  | WM only | RICOR+WM | 4thV+WM | RICOR+4thV<br>+WM | ME-ICA+WM | ME-ICA+WM<br>/<br>RICOR+4thV<br>+WM |
| --- | --- | --- | --- | --- | --- | --- |
| **GM** | 1.270<br>(0.238) | 1.353<br>(0.270) | 1.282<br>(0.241) | 1.362<br>(0.272) | 3.174<br>(0.629) | 2.368<br>(0.395) |
| **HPC** | 1.149<br>(0.256) | 1.229<br>(0.289) | 1.159<br>(0.259) | 1.237<br>(0.291) | 4.227<br>(0.856) | 3.540<br>(0.850) |
| **LC** | 1.121<br>(0.366) | 1.257<br>(0.372) | 1.141<br>(0.370) | 1.272<br>(0.372) | 3.576<br>(1.365) | 2.868<br>(0.980) |
| **MC** | 1.474<br>(0.337) | 1.559<br>(0.374) | 1.487<br>(0.341) | 1.569<br>(0.375) | 3.561<br>(0.939) | 2.312<br>(0.457) |
| **V1** | 1.231<br>(0.298) | 1.310<br>(0.328) | 1.243<br>(0.299) | 1.319<br>(0.328) | 2.554<br>(0.759) | 1.957<br>(0.429) |
| **vmPFC** | 0.533<br>(0.151) | 0.573<br>(0.173) | 0.538<br>(0.153) | 0.577<br>(0.174) | 1.923<br>(0.423) | 3.532<br>(1.010) |
| **AC** | 1.320<br>(0.306) | 1.498<br>(0.372) | 1.338<br>(0.312) | 1.513<br>(0.376) | 3.564<br>(0.952) | 2.393<br>(0.437) |

Note: Values have been multiplied by $10^3$ and calculated on data where no blurring was applied.

Analyses first focused on evaluating the effectiveness of RICOR+WM and 4thV+WM denoising on increasing tSNR in the single-echo data analyses. To do so, a repeated-measures analysis of variance (ANOVA) was performed with one random factor, participant number, and three fixed factors: ROI (7 ROI levels), application of RICOR (not applied/applied), and of 4thV regression (not included/included). tSNR varied across ROIs: it was greatest in MC and AC, and lowest in vmPFC, LC, and HPC, F(6,114) = 50.48, p < .001, η_p_^2^ = .727 (95% CI: .443 - .828). More importantly, both denoising procedures significantly increased tSNR: main effect of RICOR, F(1,19) = 91.01, p < .001, η_p_^2^ = .827 (.625 - .891), main effect of 4thV regression, F(1,19) = 142.2, p = < .001, η_p_^2^ = .882 (.736 - .925). However, their effects were under-additive, resulting in an interaction between RICOR and 4thV regression, F(1,19) = 5.67, p = .028, η_p_ = .230 (0 - .486). The effects of RICOR and 4thV regression on tSNR varied across ROIs and were largest in the MC and AC, but smallest in the vmPFC, resulting in interactions between ROI and RICOR, F(6,114) = 21.81, p < .001, η_p_ = .535 (.184 - .705), and also ROI and 4thV regression, F(6,114) = 6.377, p < .001, η_p_ = .251 (.004 - .503). Therefore, both denoising procedures removed significant but partially overlapping sources of noise from the 1E data.

A second analysis contrasted the ME-ICA-denoised data sets with the RICOR+4thV+WM-denoised 1E data. An ANOVA with denoising procedure (RICOR+4thV+WM vs. ME-ICA) and ROI as factors indicated that tSNR was significantly greater following ME-ICA denoising than following the best 1E denoising procedure, main effect of denoising procedure, F(1,19) = 279.3, p < .001, η_p_ = .936 (.853 - .959). However, the advantage was greater in some ROIs than others, denoising procedure by ROI interaction, F(6,114) = 23.85, p < .001, η_p_ = .557 (.208 - .719), main effect of ROI, F(6,114) = 37.37, p < .001, η_p_ = .663 (.344 - .788).

Relative to the best denoising procedure for 1E analyses, ME-ICA denoising increased tSNR by a factor of about 3, on average, with its largest benefits in ROIs in regions susceptible to signal dropout due to field inhomogeneities (Table 1). Because RICOR+4thV+WM produced the greatest tSNRs, subsequent analyses with 1E data were performed with this denoising pipeline.

### T2* relaxation estimates

ME-fMRI can be used to estimate T2* decay rates within each voxel to optimally combine the echoes for each voxel of the brain (Kundu et al., 2017). Variance in T2* decay rates across the brain and across individuals was examined by calculating the mean and the standard deviation of the individual T2* values in the maps generated by meica.py (Figure 3). As expected, these maps illustrate that the T2* value varies across regions: values were relatively low in vmPFC (M = 10.242, SD = 4.943), more moderate in medial occipital cortex (e.g., V1, M = 37.822, SD = 4.394), and high in the auditory cortex (M = 51.994, SD = 3.121) and motor cortex (M = 54.241, SD = 6.079). Echo times were moderate and similar for the traced LC (M = 39.360, SD = 7.241) and K1 (M = 38.059, SD = 7.928). T2* values in regions more susceptible to signal dropout due to field inhomogeneities, such as ventral temporal cortex, also showed greater variance across individuals. Therefore, at least part of the tSNR advantage for ME-ICA denoising should reflect the optimal combination of echoes using the T2* values for each individual and voxel (Kundu et al., 2017).

**Figure 3.**
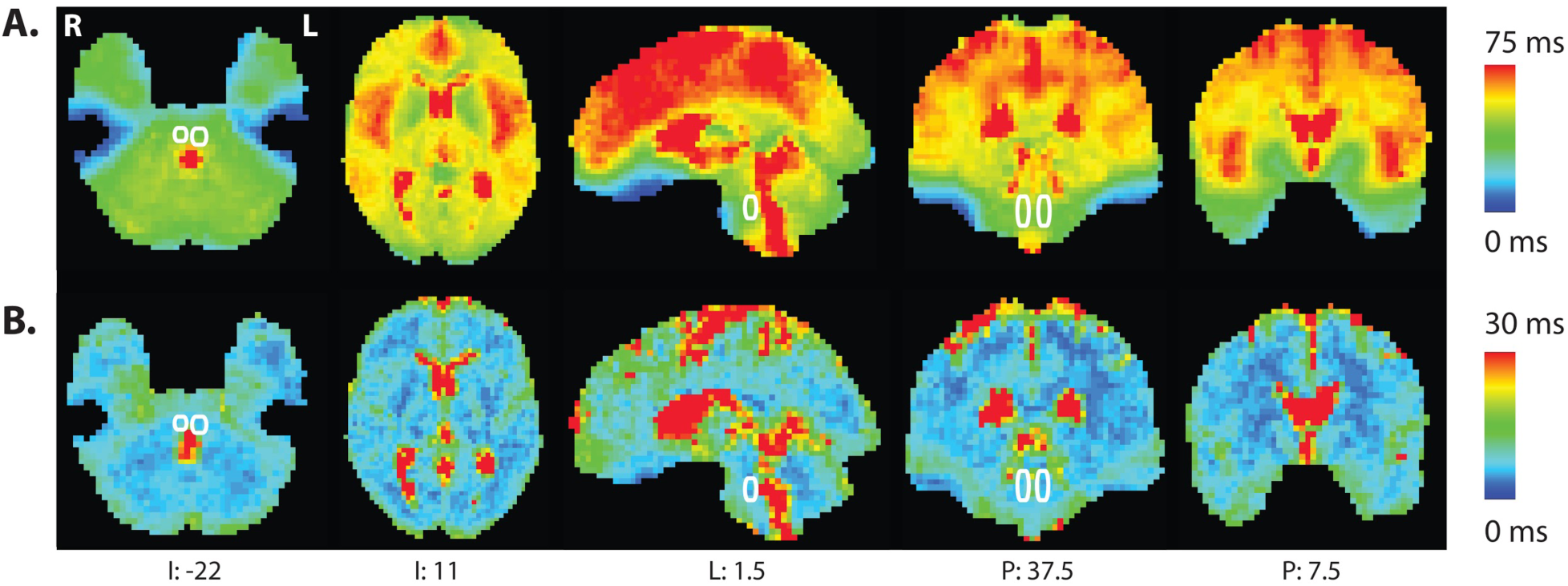
T2* map computed with t2smap.py as part of the meica.py pipeline. A. Mean observed T2* relaxation across participants. B. Standard deviation of observed T2* relaxation. The location of the LC, outlined in white, is based on the heat map illustrated in Figure 2 (coordinates in LPI).

### Intrinsic Functional Connectivity Analysis

#### Correlations Among Seeds

To examine whether estimates of LC function differ across ROI definitions, data acquisition approaches, and blurring we computed the correlation coefficients between time series of multiple brainstem regions (Table 2). Because the K1 and K2 time series ROIs were highly correlated, analyses focused on the more conservative K1 ROI.

**Table 2.** Means and standard deviations (in parentheses) of correlations among the brainstem ROI time series.

| | | $r_{K1, K2}$ | $r_{K1, PT}$ | $r_{K1, PT \cdot LC}$ | $r_{K2, PT}$ | $r_{K2, PT \cdot LC}$ | $r_{K1, LC}$ | $r_{K2, LC}$ | $r_{LC, PT}$ |
| --- | --- | --- | --- | --- | --- | --- | --- | --- | --- |
| **1E** | **No** | .860 | .399 | .384 | .470 | .458 | .125 | .136 | .158 |
|  | **blur** | (.192) | (.224) | (.224) | (.254) | (.252) | (.157) | (.183) | (.166) |
|  | **b5** | .944 | .614 | .584 | .679 | .649 | .244 | .278 | .262 |
|  |  | (.211) | (.257) | (.252) | (.278) | (.272) | (.163) | (.206) | (.189) |
| **ME** | **No** | .947 | .684 | .628 | .738 | .682 | .342 | .394 | .364 |
|  | **blur** | (.294) | (.227) | (.249) | (.249) | (.285) | (.266) | (.287) | (.282) |
|  | **b5** | .962 | .720 | .661 | .771 | .717 | .390 | .435 | .389 |
|  |  | (.275) | (.224) | (.253) | (.245) | (.286) | (.269) | (.294) | (.277) |

Note: Pearson's $r$ values were Fisher transformed to compute the mean and standard deviation and then transformed back.

Correlations between the PT and the LC and K1 ROIs were used to evaluate whether each ROI captured patterns of activity that were present in other parts of the brainstem. Correlations with the PT were greater for K1 than LC ROIs, ME than 1E data, and when the data were blurred. However, the effects of blurring were greater for 1E than ME data and for K1 than for LC. These differences were significant in an ANOVA on the Fisher-transformed correlations with PT: correlations with the PT time series significantly differed across K1 and LC ROIs, F(1,19) = 54.18, p < .001, η_p_^2^ = .740 (95% CI: .466 - .836), data type, F(1,19) = 27.06, p < .001, η_p_^2^ = .587 (95% CI: .244 - .739), and blur, F(1,19) = 231.4, p < .001, η_p_^2^ = .924 (95% CI: .826 - .952). The three way interaction was significant, F(1,19) = 24.28, p < .001, η_p_^2^ = .561 (95% CI: .213 - .722). Not surprisingly, methods that reduced noise in the data (ME-ICA denoising and blurring) increased the correlations among regions. The benefits of blurring were smaller for ME, relative to 1E, perhaps reflecting the greater tSNR (Table 1).

The moderate correlations between LC and the other ROIs, combined with the strong correlations between PT and K1 and PT and K2, suggest that the LC ROI was less likely to capture activity from the surrounding pons than were K1 and K2. To confirm this, we regressed out the LC signal from the K1, K2, and PT ROIs and correlated their residuals to produce partial correlations between K1 and PT as well as K2 and PT. These partial correlations did not differ substantially from the original correlations, indicating that the signal in the K1 and K2 ROIs that correlates with the PT reflects activity outside the individually defined LC ROIs.

### Correlations among functional connectivity maps

Differences in the iFC maps generated by each method should lead to weaker correlations among maps, particularly if those maps have already been thresholded. Therefore, the effects of ROI, ME-ICA, and blur on estimates of functional connectivity between the LC and the rest of the brain were characterized by determining the correlation and mutual information (Hausser, Strimmer, & Strimmer, 2012) of the resulting maps. For these analyses, only voxels that were in the top 15% of either map and that were not in the K1 and LC ROIs were considered because differences in high correlation voxels beyond the ROIs would likely be of most interest to researchers.

Voxel-to-voxel correlations across maps generated from 1E and ME data were small and negative, and mutual information was low (Table 3; see also Figure 4). The same was true for maps that were generated with the K1 and LC seeds (Table 4; Figure 4). Thresholded functional connectivity maps generated from 1E data rather than ME data, or from K1 rather than LC ROIs, therefore contained different information. The differences between 1E and ME maps could have resulted from the relatively large increases in tSNR when using ME data and ME-ICA, which should reduce the presence of spurious correlations in the data (Yarkoni, 2009).

**Figure 4.**
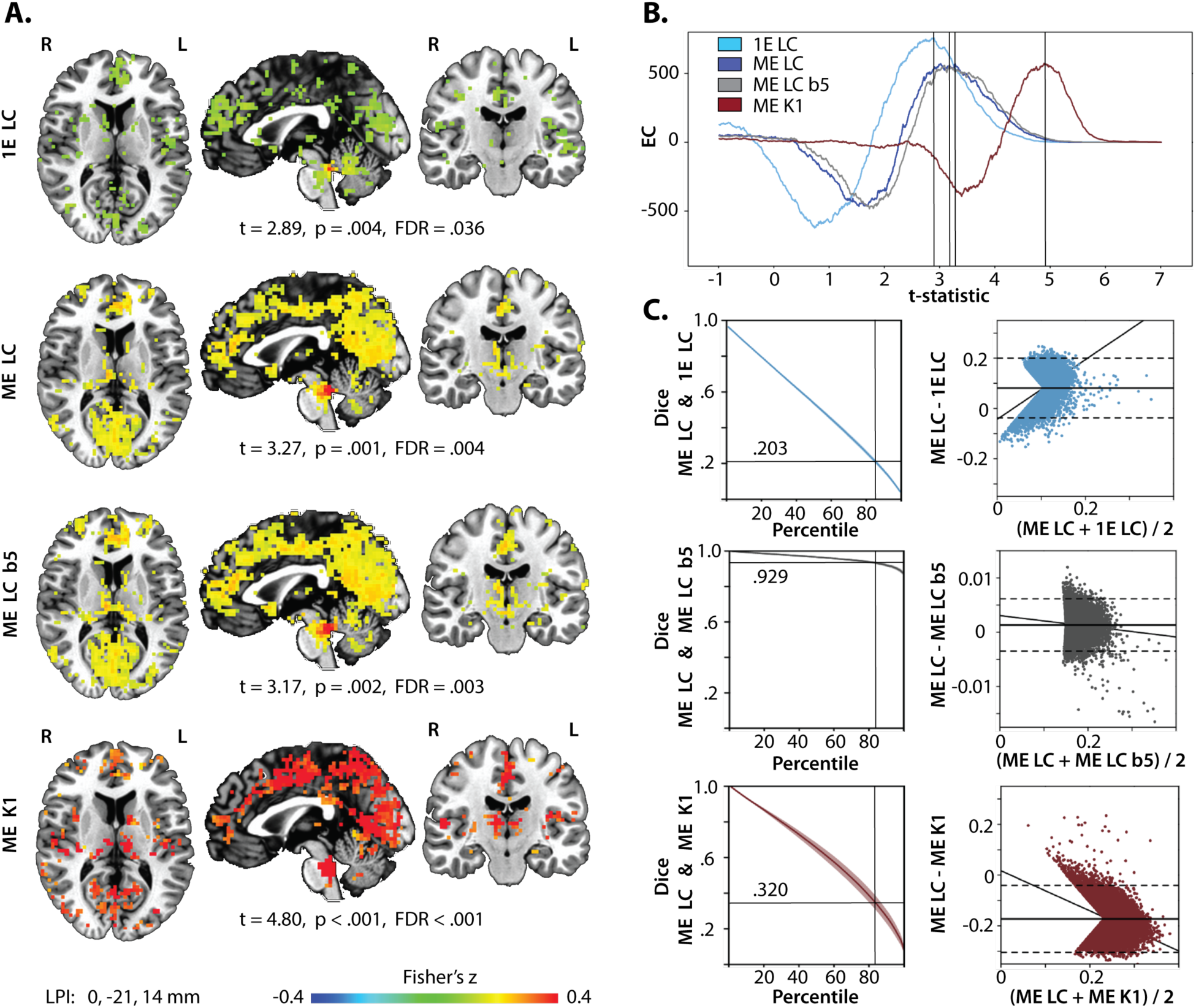
Intrinsic functional connectivity maps generated from the single-echo and multi-echo data. **A.** iFC maps for the different LC seeds with ME and 1E data. Maps were thresholded at the t-statistic with the maximum EC. B. EC for all iFC maps as a function of the t-statistic. C. Dice coefficients and Bland-Altman plots to compare the ME LC iFC map against other preprocessing approaches. Dice coefficients between the maps for all percentiles. The 85th percentile served as the cut-off for the maps used to calculate correlations and mutual information across maps reported in Tables 3-5 and to create the Bland-Altman plots in the right column. Note the smaller range on the ordinate axis for the comparison against ME LC b5. Solid line: mean difference, dashed lines: +/-1.96 of the standard deviation of the differences.

**Table 3.**
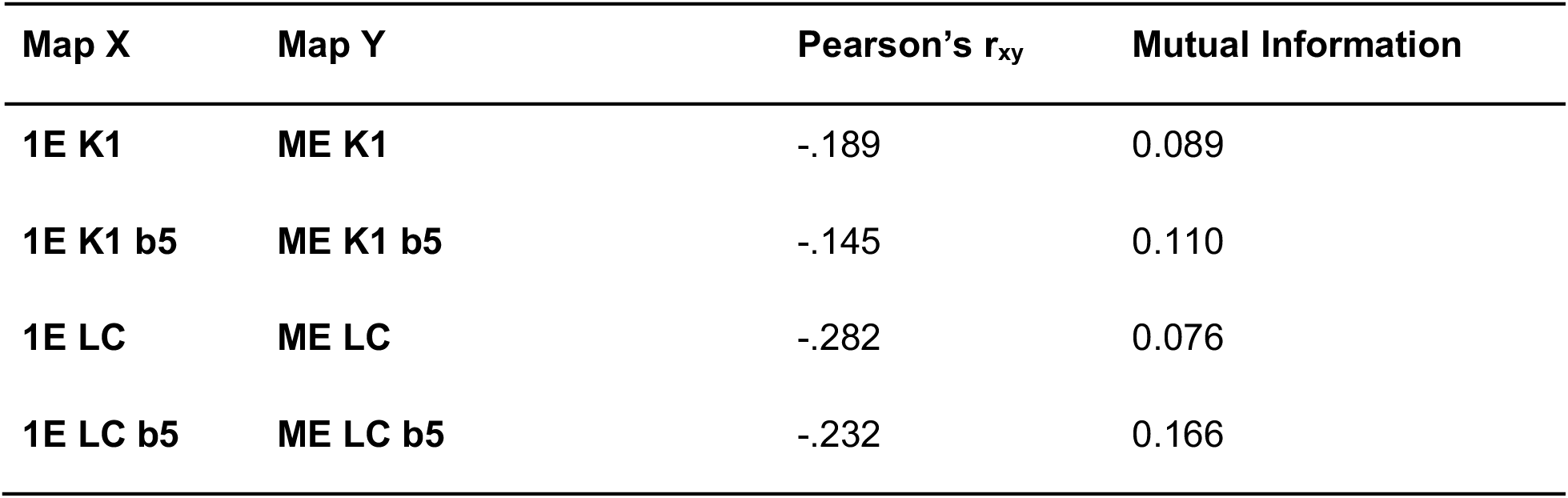
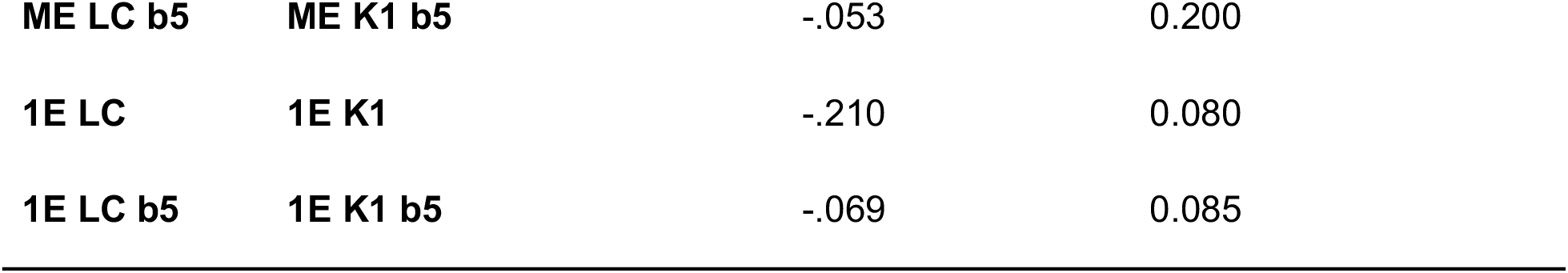
Relationships between thresholded maps that differed in data type (1E or ME)

**Table 4.**
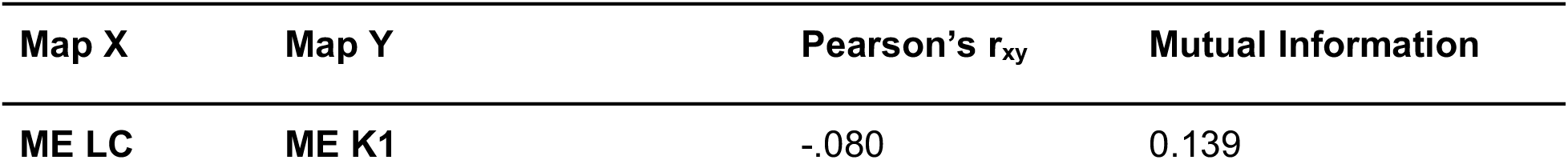
Relationships between thresholded maps that differed in seed ROI (K1 or LC)

**Table 5.** Relationships between thresholded maps that differed in blur.

| **Map X** | **Map Y** | **Pearson's $r_{xy}$** | **Mutual Information** |
| --- | --- | --- | --- |
| **ME K1** | **ME K1 b5** | .992 | 1.151 |
| **1E K1** | **1E K1 b5** | .818 | 0.421 |
| **ME LC** | **ME LC b5** | .978 | 0.556 |
| **1E LC** | **1E LC b5** | .746 | 0.199 |

However, a small amount of blurring before extracting the seed had little effect on the resulting functional connectivity maps, as indicated by the strong correlations and high amounts of mutual information (Table 5; Figure 4; also see Supplemental Information). The effects of blurring appeared to be greater for 1E data and when the LC seed was used. Thus, blurring may not produce large differences in functional connectivity maps generated from the K1 ROI, but may impact maps generated from the smaller LC ROIs.

### Characterizing the effects of seed ROI, data type, and blur on functional connectivity maps

To better characterize the extent to which different methods and preprocessing steps result in different LC connectivity maps, we visualized and quantified differences in map topography and correlation values for three central comparisons (Figure 4): (i) ME LC vs 1E LC to evaluate the effects of data type, (ii) ME LC vs ME K1 to evaluate the effects of ROI type, and (iii) ME LC vs ME LC b5 to evaluate the effects of blurring.

To quantitatively compare the topography of the functional correlation maps, the Euler characteristic (EC) was calculated with SPM12 (rev. 7487) for maps thresholded across a range of t-statistics. Briefly, negative ECs occur when the clusters are connected, but there are holes in the maps’ topography. At increasingly higher thresholds, the EC reflects the number of unconnected clusters in a given image that survive the threshold (Bowring, Maurnet, & Nichols, 2019; Brett, Penny, & Kiebel, 2003; Worsley, Marrett, Neelin, Vandal, Friston, & Evans, 1996). Therefore, each map was thresholded to the t-statistic that corresponded to the maximum EC. ECs were highest for the 1E LC map (757) and similar for the ME LC, ME LC b5, and ME K1 maps (570, 562, and 572, respectively). However, the maximum ECs occurred at lowest thresholds for the 1E LC map and at the highest threshold for the ME K1 map. The ME K1 EC was negative for the relatively high thresholds used for the other three maps, capturing widespread, nonspecific activation (particularly along the midline, see Supplementary Materials).

Agreement in which voxels survive increasingly conservative thresholds was evaluated by calculating the Dice coefficient across thresholded maps. Percentile thresholds were used to equate the number of surviving voxels in each map. Consistent with the correlation analyses, Dice coefficients decreased rapidly when the 1E LC and ME LC maps were compared (Figure 4). Decreases in the Dice coefficient for the ME LC and ME K1 comparison were less steep, but still indicated low overlap at high thresholds. Dice coefficients for the ME LC and ME LC b5 maps remained above .9 at high thresholds.

To investigate whether differences in the maps are systematic or reflect random variation across methods, Bland-Altman plots (Bland & Altman, 1999; Bowring et al., 2019) were created. These visualize differences in the estimated correlation coefficient in the two maps as a function of their mean values. Visual inspection of these plots indicate that values in the ME LC maps were systematically greater than those in the 1E LC map and systematically lower than those in the ME K1 map. In the ME K1 map, the mean of the differences fell more than 1.96 standard deviations below 0, with larger differences at more extreme mean values. Similar, but not quite as large, differences were observed when the 1E LC map was compared to the ME LC map. Blurring however, did not systematically shift the mean difference away from 0.

### Characterization of LC functional connectivity

To further characterize differences in the iFC maps, we investigated the proximity of the peaks in the ME LC map to each of the three other maps. Twenty peaks were extracted from each map using AFNI’s 3dmaxima with a minimal distance of 18 mm (6 voxels) between a given ranked peak and the next. This approach first finds the top peak in the map and then excludes any other peaks within 18 mm before searching for the next highest peak. Tables 6-9 list the top 20 peaks for all four maps as well as the rank of the nearest peak in comparison maps. Some caution is warranted in interpreting these data, since peaks that do not make it into the top 20 of one map may nevertheless fall close to a peak that did make it into the top 20 of another map.

**Table 6.** Ranked peaks in ME LC and their proximity to peaks in the other iFC maps.

| **Rank** | **Area** | **Coordinates** | **Fisher's**<br>**Z (%-tile)** | **Rank of**<br>**nearest**<br>**neighbor in**<br>**1E LC**<br>**(mm**<br>**distance)** | **Rank of**<br>**nearest**<br>**neighbor**<br>**in ME LC**<br>**b5 (mm**<br>**distance)** | **Rank of**<br>**nearest**<br>**neighbor in**<br>**ME K1 (mm**<br>**distance)** |
| --- | --- | --- | --- | --- | --- | --- |
| **1** | R. Locus<br>coeruleus | 4 -41 -19 | 0.547<br>(99.99) | 1 (6)* | 1 (0)* | 1 (15.3)* |
| **2** | L. Posterior<br>cingulate | -11 -47 8 | 0.283<br>(99.93) | 8 (9)* | 2 (0)* | 7 (4.2)* |
| **3** | R. Posterior<br>cingulate | 1 -53 23 | 0.272<br>(99.92) | 4 (25.1) | 6 (0)* | 6 (16.2)* |
| **4** | L. Anterior<br>cingulate | -11 37 17 | 0.271<br>(99.92) | 7 (11.2)* | 5 (0)* | 9 (52.5) |
| **5** | R. Anterior<br>cingulate | 7 34 23 | 0.269<br>(99.91) | 7 (9)* | 4 (0)* | 9 (45.4) |
| **6** | L. Superior Frontal gyrus | -26 31 50 | 0.266<br>(99.91) | 7 (36.2) | 3 (0)* | 9 (42.6) |
| **7** | R. Thalamus | 7 -23 14 | 0.265<br>(99.91) | 4 (17.7)* | 7 (0)* | 2 (0)* |
| **8** | L. Fusiform gyrus | -38 -59 -10 | 0.258<br>(99.89) | 6 (24.4) | 9 (9.5)* | 10 (16.4)* |
| **9** | L. Hippocampus | -17 -32 -4 | 0.254<br>(99.87) | 8 (16.4)* | 8 (3)* | 15 (7.3)* |
| **10** | L. Precuneus | -2 -53 53 | 0.253<br>(99.87) | 15 (50.9) | 16 (0)* | 4 (13.4)* |
| **11** | R. Fusiform gyrus | 31 -80 -7 | 0.252<br>(99.87) | 10 (22.6) | 13 (0)* | 5 (9.5)* |
| **12** | R. Midcingulate cortex | 4 -17 47 | 0.252<br>(99.87) | 14 (26.3) | 15 (12.4)* | 9 (15.6)* |
| **13** | R. Visual cortex / BA 17-18 | 10 -83 5 | 0.247<br>(99.84) | 15 (18)* | 17 (0)* | 5 (18)* |
| **14** | R. Cuneus | 7 -92 32 | 0.244<br>(99.81) | 15 (17.7)* | 17 (28.6) | 16 (18.2) |
| **15** | L. Middle Frontal gyrus | -29 52 23 | 0.243<br>(99.81) | 17 (27.2) | 14 (0)* | 9 (67.4) |
| **16** | R. Middle Frontal gyrus | 28 46 20 | 0.241<br>(99.87) | 17 (31.5) | 20 (0)* | 13 (56.9) |
| **17** | Subcallosal (parolfactory) area / BA 25 | 1 7 -1 | 0.240<br>(99.77) | 13 (24.2) | 7 (34.1) | 13 (24.6) |
| **18** | R. Transverse<br>temporal area<br>/ BA 42 | 46 -29 11 | 0.240<br>(99.77) | 5 (12.4)* | 12 (0)* | 8 (3)* |
| **19** | R. Precuneus | 13 -71 29 | 0.240<br>(99.77) | 15 (26.3) | 6 (22.4) | 6 (17.2)* |
| **20** | L. Thalamus | -20 -32 17 | 0.240<br>(99.77) | 8 (25.1) | 18 (0)* | 15 (18.5) |

Note: Coordinates are in LPI. \*=peaks were less than 18 mm apart and would be considered part of the same cluster by our peak finding algorithm.

**Table 7.** Ranked peaks in the 1E LC iFC map.

| **Rank** | **Area** | **Coordinates** | **Fisher's Z (%-tile)** | **Rank of nearest neighbor in ME LC (mm)** |
| --- | --- | --- | --- | --- |
| **1** | R. Locus Coeruleus* | -2 -41 -19 | 0.405 (99.99) | 1 (6) |
| **2** | L. Cerebellum | -26 -83 -22 | 0.168 (99.88) | 8 (29.4) |
| **3** | R. Cerebellum | 1 -59 -19 | 0.166 (99.87) | 1 (18.2) |
| **4** | R. Hippocampus* | 10 -38 5 | 0.163 (99.86) | 7 (17.7) |
| **5** | R. Posterior Insula* | 46 -32 23 | 0.153 (99.82) | 18 (12.4) |
| **6** | L. Cerebellum | -23 -44 -22 | 0.152 (99.82) | 9 (22.4) |
| **7** | L. Anterior Cingulate* | -2 34 23 | 0.148 (99.80) | 5 (9) |
| **8** | L. Posterior Cingulate* | -11 -47 -1 | 0.144 (99.78) | 2 (9) |
| **9** | L. Anterior Insula | -47 7 -1 | 0.137 (99.71) | 17 (48) |
| **10** | R. Cerebellum | 19 -65 -19 | 0.136 (99.71) | 11 (22.6) |
| **11** | R. Anterior Insula | 40 7 -10 | 0.136 (99.71) | 17 (40) |
| **12** | L. Medial Frontal gyrus | -2 52 -4 | 0.132 (99.65) | 4 (27.3) |
| **13** | R. Ventral Tegmental Area | 1 -17 -4 | 0.130 (99.64) | 7 (19.9) |
| **14** | R. Superior Frontal gyrus | 10 7 56 | 0.130 (99.64) | 12 (26.3) |
| **15** | L. Visual cortex / BA 17-18* | -2 -89 17 | 0.130 (99.91) | 14 (17.7) |
| **16** | L. Superior Temporal gyrus | -53 -17 -4 | 0.129 (99.62) | 9 (39) |
| **17** | L. Medial Frontal gyrus | -2 55 23 | 0.129 (99.61) | 4 (21) |
| **18** | L. Visual cortex / V1 | -5 -98 -1 | 0.128 (99.58) | 13 (22) |
| **19** | R. Lateral Occipital area | 40 -74 14 | 0.127 (99.57) | 11 (23.6) |
| **20** | R. Precentral gyrus | 40 -20 44 | 0.126 (99.54) | 18 (34.7) |

Note: \*=peak fell within 18 mm of a peak in the ME LC map and would be considered part of the same cluster by our algorithm.

**Table 8.** Ranked peaks in the ME LC b5 iFC map.

| **Rank** | **Area** | **Coordinates** | **Fisher's Z (%-tile)** | **Rank of nearest neighbor in ME LC (mm)** |
| --- | --- | --- | --- | --- |
| **1** | R. Locus Coeruleus* | 4 -41 -19 | 0.559 (99.99) | 1 (0) |
| **2** | L. Posterior cingulate* | -11 -47 8 | 0.295 (99.93) | 2 (0) |
| **3** | L. Superior Frontal gyrus* | -26 31 50 | 0.286 (99.91) | 6 (0) |
| **4** | R. Anterior cingulate* | 7 34 23 | 0.280 (99.91) | 5 (0) |
| **5** | L. Anterior cingulate* | -11 37 17 | 0.279 (99.90) | 4 (0) |
| **6** | R. Posterior cingulate* | 1 -53 23 | 0.276 (99.90) | 3 (0) |
| **7** | R. Thalamus* | 7 -23 14 | 0.275 (99.90) | 7 (0) |
| **8** | L. Hippocampus* | -20 -32 -4 | 0.273 (99.90) | 9 (3) |
| **9** | L. Fusiform gyrus* | -29 -56 -10 | 0.270 (99.89) | 8 (9.5) |
| **10** | R. Hippocampus | 16 -35 -1 | 0.264 (99.87) | 7 (21.2) |
| **11** | L. Midcingulate cortex | -2 -32 50 | 0.264 (99.87) | 12 (16.4) |
| **12** | R. Transverse temporal<br>area / BA 42* | 46 -29 11 | 0.257 (99.85) | 18 (0) |
| **13** | R. Fusiform gyrus* | 31 -80 -7 | 0.256 (99.84) | 11 (0) |
| **14** | L. Middle Frontal gyrus* | -29 52 23 | 0.255 (99.83) | 15 (0) |
| **15** | R. Midcingulate gyrus* | 1 -5 47 | 0.255 (99.83) | 12 (12.4) |
| **16** | L. Precuneus* | -2 -53 53 | 0.255 (99.83) | 10 (0) |
| **17** | R. Visual cortex / BA 17-18* | 10 -83 5 | 0.253 (99.81) | 13 (0) |
| **18** | L. Thalamus* | -20 -32 17 | 0.250 (99.78) | 20 (0) |
| **19** | L. Anterior Insula | -35 10 -7 | 0.249 (99.77) | 17 (36.6) |
| **20** | R. Middle Frontal gyrus* | 28 46 20 | 0.249 (99.77) | 16 (0) |

Note: \*=peak fell within 18 mm of a peak in the ME LC map and would be considered part of the same cluster by our algorithm.

**Table 9.** Ranked peaks in the ME K1 iFC map.

| **Rank** | **Area** | **Coordinates** | **Fisher's Z (%-**<br>**tile)** | **Rank of nearest**<br>**neighbor in ME**<br>**LC (mm)** |
| --- | --- | --- | --- | --- |
| **1** | L. Locus Coeruleus* | -5 -38 -31 | 0.830 (99.99) | 1 (15.3) |
| **2** | R. Thalamus* | 7 -23 14 | 0.495 (99.90) | 7 (0) |
| **3** | R. Hippocampus | 19 -32 -4 | 0.487 (99.89) | 1 (23) |
| **4** | L. Precuneus* | -2 -65 59 | 0.482 (99.89) | 10 (13.4) |
| **5** | R. Lingual gyrus* | 22 -77 -7 | 0.480 (99.88) | 11 (9.5) |
| **6** | R. Calcarine gyrus* | 1 -68 17 | 0.479 (99.88) | 3 (16.2) |
| **7** | L. Posterior cingulate* | -8 -47 11 | 0.475 (99.88) | 2 (4.2) |
| **8** | R. Transverse temporal<br>area / BA 42* | 46 -29 14 | 0.475 (99.87) | 18 (3) |
| **9** | R. Midcingulate gyrus* | 1 -2 50 | 0.469 (99.86) | 12 (15.6) |
| **10** | L. Fusiform gyrus* | -53 -65 -13 | 0.465 (99.84) | 8 (16.4) |
| **11** | R. Fusiform gyrus | 40 -65 -16 | 0.463 (99.84) | 11 (19.7) |
| **12** | L. Midcingulate gyrus | -5 -32 50 | 0.462 (99.83) | 12 (17.7) |
| **13** | R. Hippocampus | 22 -2 -10 | 0.459 (99.82) | 17 (24.6) |
| **14** | R. Visual cortex / BA 17-18 | 13 -101 -4 | 0.455 (99.80) | 13 (20.3) |
| **15** | L. Hippocampus* | -23 -35 -1 | 0.451 (99.76) | 9 (7.3) |
| **16** | R. Cuneus | 25 -89 32 | 0.450 (99.76) | 14 (18.2) |
| **17** | L. Posterior Insula | -47 -26 17 | 0.450 (99.76) | 20 (27.7) |
| **18** | L. Precuneus | -11 -59 74 | 0.443 (99.69) | 10 (23.6) |
| **19** | R. Supplementary Motor<br>Area | 1 -8 74 | 0.442 (99.68) | 12 (28.6) |
| **20** | L. Lateral Occipital Area | -47 -83 5 | 0.442 (99.67) | 8 (29.7) |

Note: \*=peak fell within 18 mm of a peak in the ME LC map and would be considered part of the same cluster by our algorithm.

Consistent with the previous analyses, the percentage of peaks in the ME LC map that fell within 18 mm of a peak on another map was highest for the ME LC b5 map (17 of 20 peaks, or 85%), next highest for ME K1 map (12 of 20 peaks, or 60%), and lowest for the 1E LC maps (9 of 40 peaks, or 45%).

## General Discussion

A growing body of research on the functional and anatomical properties of the LC suggest that it plays a central role in a variety of cognitive processes and in neurodegenerative disease (Mather & Harley, 2016). However, characterizing LC function and anatomy in humans using fMRI has multiple challenges that have not been systematically addressed in the literature (Liu et al., 2017). This study addressed several of these challenges by comparing LC localization approaches, BOLD data acquisition and denoising approaches, and the use of conservative blurring on estimates of LC activity and functional connectivity. The analyses suggest that, while a small amount of blurring may not have a large effect on estimates of LC function, LC localization, data acquisition, and denoising approaches may lead to different results.

### Localization of LC with a probabilistic atlas or nmT1 imaging

One purpose of this study was to investigate whether individually defining LC ROIs using nmT1 imaging produces different results than using a probabilistic atlas. Probabilistic ROIs may not adequately account for variability in individual brainstem anatomy and LC shape and location (German et al., 1988; Keren et al., 2015). In addition, studies report a wide range of bilateral LC coordinates that sometimes go beyond the dimensions of the Keren atlas (Liu, et al., 2017). The use of an ROI defined by coordinates or a group atlas, like the K1 ROIs used here, could therefore miss the LC entirely, include only part of the LC, and capture activity from neighboring brainstem areas (e.g., the pontine tegmentum, medial cerebral peduncle, cranial nerve nuclei, and inferior colliculus; Neary, 2008). These neighboring areas play critical roles in processing motor and sensory information, which could confound analyses. Thus, the best way to isolate activity from the LC could still be to individually define ROIs using nmT1 imaging.

The data supported this recommendation: the K1 ROIs were larger and tended to extend further rostrally and caudally than the individual LC ROIs, and there was only low to moderate overlap between the individually defined LC ROIs and K1 ROIs. These differences had consequences for estimates of LC connectivity. The correlation between K1 and the LC ROI was weaker than the correlation between K1 and a different, non-overlapping region of the pons, the pontine tegmentum. In addition, agreement between thresholded iFC maps generated by the LC and K1 ROIs was low (.3 for the top 15% of voxels). iFC maps created with the Keren atlas showed stronger correlations than the LC overall, which caused less specific activity at thresholds used for the other maps, and the need for higher thresholds to isolate network clusters. Both the LC and K1 ROIs produced maps with peaks that were near each other, including thalamus, HPC, areas along the midline, and sensory areas. However, peaks that did not appear to have a match in the other map were observed in precentral gyrus, middle frontal gyrus, anterior cingulate, and anterior insula. Thus, the final iFC maps produced by K1 showed more widespread associations that also differed in their spatial topography from those produced by the nmT1-based LC ROIs.

### Use of ME-fMRI or 1E-MRI

Because ME-fMRI with ME-ICA denoising may be particularly effective at reducing noise from motion and improving BOLD contrast (Power et al., 2018), another primary goal of this study was to characterize whether the resting state iFC maps differed when ME data rather than 1E data were analyzed. Analyses suggested that they do: overlap between the two maps decreased rapidly as thresholds increased, correlations between the time series and group level iFC maps of the 1E and ME LC data sets were low, and only 45% of the peaks identified by either map were within 18 mm of a peak in the other map. Unlike the ME LC map, the 1E LC map had many highly ranked peaks in the cerebellum. These differences likely reflect the ability to accommodate variance in the optimal echo time across voxels and to remove components of the signal with non-BOLD-like characteristics (Kundu et al., 2012). Indeed, as demonstrated by the three-fold increase in tSNR, ME-fMRI with ME-ICA was superior to 1E-fMRI that used respiration and cardiac measures for cleaning the data (RETROICOR; both included WM and 4thV nuisance regressors). This improvement in tSNR was evident in every region examined, including the LC ROIs. Thus, the use of ME-fMRI appears to be advantageous, particularly with small regions like the LC.

Although ME-ICA is a relatively effective way to remove some sources of noise from fMRI data, it may be necessary to employ additional denoising methods to remove respiratory and cardiac noise in the BOLD signal itself (Power et al., 2018). Our ME-fMRI pipeline therefore included both WM and 4thV regression (Dagli, Ingeholm, & Haxby, 1999; Windischberger et al., 2002). However, analyses that did not include the WM nuisance regressor resulted in similar outcomes, with correlations that were somewhat larger (Supplemental Materials).

### Effects of blurring on the data

Because of the LCs size and proximity to the fourth ventricle, this study also addressed the effects of blurring prior to extracting the seed from the ROIs. The results suggest that this had only a small effect on the resulting iFC maps: correlations among the seeds and resulting maps were strong, ECs for the two maps were similar, Dice coefficients were high, and Bland-Altman plots suggested small differences in the absolute magnitude of the maps. This resulted in 70% of the peaks in both maps being within 18 mm of a cluster in the other map. Peaks that were further away were also in relatively similar brain regions. This suggests that blurring by a small amount before the LC seed is extracted may not have an appreciable effect on the conclusions of a study. Blurring by larger amounts, however, may pose bigger problems, as it is likely to increase the similarity between the LC and surrounding pontine areas. As the K1 ROI analyses demonstrate, this could lead to differential outcomes in the data (see also Supplemental Materials for maps with blurring to 8 mm FWHM).

### Measuring neuromelanin content with nmT1 imaging

Although not the focus of the current study, many studies of the LC are concerned with individual differences in neuromelanin content. Liu, et al. (2017) identified substantial methodological differences in how neuromelanin content is estimated using nmT1 images. Studies that use nmT1 signal intensity to estimate LC neuromelanin content have not always normalized signal intensity (Liu, et al., 2017) and those studies that did often used different reference regions. Among the studies that normalized signal intensity, the most common reference region was the PT, a large region with vague boundaries. Because the PT displays age-related signal decline, including in non-clinical populations, (Clewett et al., 2016), it may not be an optimal choice for a reference. Studies also differ in the algorithms used to normalize signal intensity, differing in their use of peak or average LC signal. One consequence of these methodological differences may be greater variability in estimates of LC volume, which vary widely in reported studies and are different than what one would expect from post-mortem specimens (Liu, et al., 2017). Despite these problems, nmT1 imaging remains the best avenue to measure in vivo individual differences in signal intensity. To mitigate some of these issues in our preprocessing of the nmT1 images, we applied a novel normalization approach that does not rely on a specific reference region, but instead relies on variance within the brainstem itself. This circumvents some of the problems with using a reference region.

### Functional connectivity of the LC in typical young adults

Previous functional studies on the LC have exclusively been done using 1E-fMRI and frequently use the Keren 2 SD atlas. Across those studies, regions that typically emerge in iFC analyses are the prefrontal cortex, cerebellum, right supplementary motor area, cingulate cortex, thalamus, visual cortex, lingual and fusiform gyrus, insula, amygdala, and the hippocampus (e.g., Bär et al., 2016; Köhler, Wagner, & Bär, 2016; Kline et al., 2016; Murphy et al., 2014; Sterpenich et al., 2006). Nearly all these regions were ranked highly across the 1E LC, ME LC, and ME K1 maps. Only a few were not represented in every pipeline. However, the amygdala did not rank in the top 20 of any of our maps, possibly because it was too close to peaks in the hippocampus to survive our peak selection procedure.

Previous studies on LC iFC do not always identify the same regions. For example, Song et al. (2017) showed connectivity to the reticular formation, ventral tegmental area, and caudate. None of our pipelines ranked peaks in the reticular formation or caudate highly, although these (or nearby) regions did survive our Euler characteristic-based thresholding procedure. The ventral tegmental area was ranked 13th in the 1E LC map. Zhang, Hu, Chao, & Li (2016) found positive connectivity to inferior temporal cortex, anterior parahippocampal gyrus, posterior insula, ventrolateral thalamus, and a large region in the cerebellum. Intriguingly, they found negative connectivity to several regions that were positively correlated with the LC in our maps (similar to others), including the bilateral visual cortex, middle and superior temporal cortex, precuneus, retrosplenial cortex, posterior parahippocampal cortex, frontopolar cortex, caudate, and dorsal and medial thalamus. An important methodological difference between our study and theirs, however, was the use of whole brain (global) signal regression, which can introduce negative correlations (Fox, Zhang, Snyder, & Raichle, 2009). We will return to this point later.

One intriguing difference between our ME LC maps and the 1E LC or ME K1 maps was the presence of a cluster in the basal forebrain (labeled subcallosal area in Table 4a). This cluster has not been widely reported in studies of LC iFC, perhaps because it is located in a part of the brain that is susceptible to field inhomogeneities and difficult to measure using 1E-fMRI. Previous work has demonstrated that ME-fMRI can effectively recover signal from this region during pre-processing (Markello, et al., 2018). The presence of this region in our ME LC maps is particularly important, however, as studies on the basal forebrain’s physiology and function suggest an important anatomical and functional relationship between these regions and the LC in the regulation of attention and memory (e.g., Mesulam, 2013; Ljubojevic, Luu, & De Rosa, 2014; Sarter, Hasselmo, Bruno, & Givens, 2005; Yu & Dayan, 2005)

### Pupillometry as a means of investigating LC function

Pupillometry is increasingly used as a non-invasive proxy measure for LC activity (e.g., Einhäuser, Stout, Koch, & Carter, 2008; Eldar, Niv, & Cohen, 2013; Gilzenrat, Nieuwenhuis, Jepma, & Cohen, 2010; Jepma & Nieuwenhuis, 2011; Murphy et al., 2014). Recent neuroimaging and electrophysiological work has shown a strong, positive relationship between activity in the LC and increases in pupil size (Breton-Provencher & Sur, 2019; Joshi, Li, Kalwani, & Gold, 2016; Murphy et al., 2014). However, they also demonstrate that pupil diameter is associated with activity in other brainstem areas, including those that are near the LC (e.g., inferior colliculus, superior colliculus, nucleus incertus; e.g., Costa & Rudebeck, 2016; Joshi et al., 2016; Szőnyi et al., 2019). Efforts to use pupillometry to verify that certain voxels are within the LC have found activations throughout the brainstem, particularly in dorsal areas (near the colliculi), and particularly in oddball tasks (Murphy et al., 2014). This, combined with our data, suggest caution is warranted when using pupillometry and K1 ROIs to isolate LC activity: larger ROIs from probabilistic atlases, like the K1 ROIs, produce appreciably different iFC maps than the individual LC ROIs.

### Limitations and considerations

There are several limitations to the results presented here. First, we did not explore other denoising approaches for 1E or ME data (e.g., ANATICOR and ICA AROMA) and it is possible that these would have led to different outcomes (e.g., Dipasquale et al., 2017; Jo, Saad, Simmons, Milbury, & Cox, 2010; for a review of available denoising procedures, see Caballero-Gaudes & Reynolds, 2017). Second, a common denoising strategy in resting state iFC analyses is to regress out the mean signal across the brain (global signal) to reduce the contributions of physiological and motion related artifacts (Liu, Nalci, & Falahpour, 2017). However, the global signal contains functional signals (e.g., Schölvinck et al., 2010), including those related to vigilance, that would be removed if global signal regression were employed (e.g., Liu et al., 2017). This is of particular concern for studies of the LC because it is involved in regulating vigilance both on and off-task (e.g., Aston-Jones et al., 1994; Aston-Jones et al., 2007), and has widespread projections throughout the brain (e.g., Schwarz & Luo, 2015). To account for physiological noise in the data all pre-processing approaches in this study included WM and 4thV nuisance regressors. Although we did not systematically explore the effects of global signal regression on the outcome of LC iFC analyses, versions of the ME LC and 1E LC maps with global signal regression rather than WM and 4thV nuisance regressors are in the Supplementary Materials. Correlation coefficients were weaker in these maps and included negative values (a common consequence of global signal regression; Fox et al., 2009). Pontine areas centered on the respective seed were the only region across all four pipelines that survived global signal regression. A cluster in the medial prefrontal region was also still present in all ME-fMRI maps but not the 1E-fMRI map,. The ME K1 map also included a cluster in the thalamus. Zhang et al. (2016) applied global signal regression to their data and found negative connectivity with, amongst other areas, visual cortex. We did not see that pattern in our data. However, global signal regression did introduce negative correlations diffusely spread throughout lateral temporal cortex, around the ventricles, and the cerebellum.

There are also pragmatic concerns associated with ME-fMRI, which requires longer TRs than 1E-fMRI and greater computational resources. Shorter TRs are sometimes needed to characterize the hemodynamic response or when tasks include events that are closely spaced. Under these conditions, multi-echo multi-band techniques (MEMB; e.g., Preibish, Castrillón G, Bührer, & Riedl, 2015) could be used. One systematic evaluation of ME-ICA with MEMB suggests that this approach may offer additional benefits by improving estimates of the hemodynamic response (Kundu et al., 2017). However, ME-fMRI also has higher computational costs than 1E-fMRI that would be exacerbated by combining it with multi-band imaging. Because acquiring three echoes triples the amount of data acquired during a scan, ME-fMRI requires more storage than 1E-fMRI. In addition, running ME-ICA adds substantial processing time to preprocessing. Depending on a given study’s experimental goals, timeframe, and budget, these factors may reduce the utility of ME-fMRI with ME-ICA, despite the clear and well-established boost to data quality and power.

The ability to use nmT1 imaging may also be limited to particular scanners and populations. Neuromelanin levels change over the lifespan (Halliday, Fedorow, Rickert, Gerlach, Riederer, & Double, 2006) and differ between clinical and non-clinical populations (Braak, Thal, Ghebremedhin, & Del Tredici, 2011; Marien, Colpaert, & Rosenquist, 2004). Researchers interested in the effects of neuromelanin on cognitive decline in the elderly, cognitive development in childhood, or the mediating role of neuromelanin on cognitive differences between clinical and non-clinical populations, may therefore have to consider alternative ways of localizing the LC. Furthermore, nmT1 scanning works best with 3T scanners, as it has a better signal to noise ratio than 1.5T (Uğurbil, Garwood, & Ellermann, 1993) but none of the unique challenges that come with human imaging at 7T or higher (Ladd et al., 2018).

Even when the LC can be localized with nmT1 imaging, other difficulties emerge. First, many registration algorithms are optimized for aligning the neocortex rather than the brainstem to an atlas. The brainstem may be best aligned to group data if it is separated from the rest of the brain and nonlinearly warped to the template (Tona et al., 2017) and such algorithms are not widely used. Second, the small size of the LC and coarse spatial resolution of functional images make it likely that aligning individually defined LC ROIs to an atlas may distort the ROI (binary masks become proportional following alignment and must be thresholded again), causing the resulting ROI to include signal from nearby pontine structures. This is particularly true for studies, like ours, that use 3 mm isotropic voxels (within-plane diameter of the LC is about 2.5 mm; Fernandes et al., 2012).

## Conclusion

fMRI studies of LC function and structure have been hampered by the difficulties involved in localizing the LC, isolating LC activity from other nearby regions, and removing the effects of noise from motion and physiology on BOLD data (Liu, et al., 2017). By systematically investigating the benefits of using nmT1 images, ME-fMRI, and conservative amounts of blurring, this study has found that the methods used to localize the LC, acquire BOLD data, and denoise it may have significant effects on how LC function is characterized. Although using nmT1 images to localize the LC necessitates additional scan time and manual tracing, these efforts lead to greater specificity in the iFC maps. In addition, ME-fMRI with ME-ICA denoising protocols increased the quality of the data and revealed a cluster in the basal forebrain, a region that is otherwise susceptible to signal drop out (Park et al., 1988).

## Author Acknowledgments

The authors would like to thank Emily Qualls, Roy Proper, Alyssa Phelps, Bohan Li, and Roy Moyal for their assistance with data collection. Thanks also to Adam Anderson and Eve De Rosa for their input on this work. We also thank Mark Eckert for providing the LC probabilistic atlas.

## Abbreviations

LC: locus coeruleus
4thV: 4th ventricle
NE: norepinephrine
nmT1: neuromelanin-weighted
T1 iFC: intrinsic functional connectivity
1E-fMRI: single-echo fMRI
ME-fMRI: multi-echo fMRI
K1, K2: binary atlas from Keren, Lozar, Harris, Morgan, & Eckert (2009), 1SD and 2SD

## References

Astafiev, S. V., Snyder, A. Z., Shulman, G. L., & Corbetta, M. (2010). Comment on “Modafinil shifts human locus coeruleus to low-tonic, high-phasic activity during functional MRI” and “Homeostatic sleep pressure and responses to sustained attention in the suprachiasmatic area.” Science, 328(5976), 309–309. https://doi.org/10.1126/science.1177200

Aston-Jones, G., Shipley, M. T., Chouvet, G., Ennis, M., van Bockstaele, E., Pieribone, V., … Williams, J. T. (1991). Chapter 4 - Afferent regulation of locus coeruleus neurons: Anatomy, physiology and pharmacology. In C. D. Barnes & O. Pompeiano (Eds.), Progress in Brain Research, (Vol. 88, pp. 47–75). Elsevier Press. https://doi.org/10.1016/S0079-6123(08)63799-1

Aston-Jones, Gary, Gonzalez, M., & Doran, S. (2007). Role of the locus coeruleus-norepinephrine system in arousal and circadian regulation of the sleep-wake cycle. In Brain norepinephrine: Neurobiology and therapeutics(pp. 157–195). https://doi.org/10.1017/CBO9780511544156.007

Aston-Jones, Gary, Rajkowski, J., & Cohen, J. (1999). Role of locus coeruleus in attention and behavioral flexibility. Biological Psychiatry, 46(9), 1309–1320. https://doi.org/10.1016/S0006-3223(99)00140-7

Aston-Jones, Gary, & Waterhouse, B. (2016). Locus Coeruleus: From Global Projection System to Adaptive Regulation of Behavior. Brain Research, 1645, 75–78. https://doi.org/10.1016/j.brainres.2016.03.001

Bär, K.-J., de la Cruz, F., Schumann, A., Koehler, S., Sauer, H., Critchley, H., & Wagner, G. (2016). Functional connectivity and network analysis of midbrain and brainstem nuclei. NeuroImage, 134, 53–63. https://doi.org/10.1016/j.neuroimage.2016.03.071

Berridge, C. W., & Waterhouse, B. D. (2003). The locus coeruleus–noradrenergic system: Modulation of behavioral state and state-dependent cognitive processes. Brain Research Reviews, 42(1), 33–84. https://doi.org/10.1016/S0165-0173(03)00143-7

Biswal, B., Yetkin, F. Z., Haughton, V. M., & Hyde, J. S. (1995). Functional connectivity in the motor cortex of resting human brain using echo-planar mri. Magnetic Resonance in Medicine, 34(4), 537–541. https://doi.org/10.1002/mrm.1910340409

Bland, J. M., & Altman, D. G. (1999). Measuring agreement in method comparison studies. Statistical Methods in Medical Research, 8(2), 135–160. https://doi.org/10.1177/096228029900800204

Bowring, A., Maumet, C., & Nichols, T. E. (2019). Exploring the impact of analysis software on task fMRI results. Human Brain Mapping, 40(11), 3362–3384. https://doi.org/10.1002/hbm.24603

Braak, H., Thal, D. R., Ghebremedhin, E., & Del Tredici, K. (2011). Stages of the Pathologic Process in Alzheimer Disease: Age Categories From 1 to 100 Years. Journal of Neuropathology & Experimental Neurology, 70(11), 960–969. https://doi.org/10.1097/NEN.0b013e318232a379

Brett, M., Penny, W., & Kiebel, S. (2003). *An Introduction to Random Field Theory*. In R. Frackowiak, K. Friston, C. Frith, R. Dolan, C. Price, S. Zeki, J. Ashburner, & W. Penny (Eds.), Human Brain Function II, (Chapter 14). Academic Press.

Breton-Provencher, V., & Sur, M. (2019). Active control of arousal by a locus coeruleus GABAergic circuit. Nature Neuroscience, 22(2), 218–228. doi:10.1038/s41593-018-0305-z

Chandler, D. J., Gao, W.-J., & Waterhouse, B. D. (2014). Heterogeneous organization of the locus coeruleus projections to prefrontal and motor cortices. Proceedings of the National Academy of Sciences of the United States of America, 111(18), 6816–6821. https://doi.org/10.1073/pnas.1320827111

Cho, Z. H., & Ro, Y. M. (1992). Reduction of susceptibility artifact in gradient-echo imaging. Magnetic Resonance in Medicine, 23(1), 193–200. https://doi.org/10.1002/mrm.1910230120

Clewett, D. V., Lee, T.-H., Greening, S., Ponzio, A., Margalit, E., & Mather, M. (2016). Neuromelanin marks the spot: Identifying a locus coeruleus biomarker of cognitive reserve in healthy aging. Neurobiology of Aging, 37, 117–126. https://doi.org/10.1016/j.neurobiolaging.2015.09.019

Costa, V. D., & Rudebeck, P. H. (2016). More than Meets the Eye: The Relationship between Pupil Size and Locus Coeruleus Activity. Neuron, 89(1), 8–10. https://doi.org/10.1016/j.neuron.2015.12.031

Cox, R. W. (1996). AFNI: Software for Analysis and Visualization of Functional Magnetic Resonance Neuroimages. Computers and Biomedical Research, 29(3), 162–173. https://doi.org/10.1006/cbmr.1996.0014

Cox, R. W., & Jesmanowicz, A. (1999). Real-time 3D image registration for functional MRI. Magnetic Resonance in Medicine, 42(6), 1014–1018. https://doi.org/10.1002/(SICI)1522-2594(199912)42:6<1014::AID-MRM4>3.0.CO;2-F

Dagli, M. S., Ingeholm, J. E., & Haxby, J. V. (1999). Localization of cardiac-induced signal change in fMRI. NeuroImage, 9(4), 407–415. https://doi.org/10.1006/nimg.1998.0424

Dale, A. M., Fischl, B., & Sereno, M. I. (1999). Cortical Surface-Based Analysis: I. Segmentation and Surface Reconstruction. NeuroImage, 9(2), 179–194. https://doi.org/10.1006/nimg.1998.0395

Dipasquale, O., Sethi, A., Laganà, M. M., Baglio, F., Baselli, G., Kundu, P., … Cercignani, M. (2017). Comparing resting state fMRI de-noising approaches using multi- and single-echo acquisitions. PLoS ONE, 12(3), e0173289. https://doi.org/10.1371/journal.pone.0173289

Einhauser, W., Stout, J., Koch, C., & Carter, O. (2008). Pupil dilation reflects perceptual selection and predicts subsequent stability in perceptual rivalry. Proceedings of the National Academy of Sciences, 105(5), 1704–1709. https://doi.org/10.1073/pnas.0707727105

Eldar, E., Niv, Y., & Cohen, J. D. (2016). Do You See the Forest or the Tree? Neural Gain and Breadth Versus Focus in Perceptual Processing. Psychological Science, 27(12), 1632–1643. https://doi.org/10.1177/0956797616665578

Enochs, W. S., Petherick, P., Bogdanova, A., Mohr, U., & Weissleder, R. (1997). Paramagnetic metal scavenging by melanin: MR imaging. Radiology, 204(2), 417–423. https://doi.org/10.1148/radiology.204.2.9240529

Fernandes, P., Regala, J., Correia, F., & Gonçalves-Ferreira, A. J. (2012). The human locus coeruleus 3-D stereotactic anatomy. Surgical and Radiologic Anatomy, 34(10), 879–885. https://doi.org/10.1007/s00276-012-0979-y

Fischl, B., Sereno, M. I., & Dale, A. M. (1999). Cortical Surface-Based Analysis: II: Inflation, Flattening, and a Surface-Based Coordinate System. NeuroImage, 9(2), 195–207. https://doi.org/10.1006/nimg.1998.0396

Foote, S. L., Freedman, R., & Oliver, A. P. (1975). Effects of putative neurotransmitters on neuronal activity in monkey auditory cortex. Brain Research, 86(2), 229–242. https://doi.org/10.1016/0006-8993(75)90699-X

Fox, M. D., Zhang, D., Snyder, A. Z., & Raichle, M. E. (2009). The Global Signal and Observed Anticorrelated Resting State Brain Networks. Journal of Neurophysiology, 101(6), 3270–3283. https://doi.org/10.1152/jn.90777.2008

Genovese, C. R., Lazar, N. A., & Nichols, T. (2002). Thresholding of statistical maps in functional neuroimaging using the false discovery rate. NeuroImage, 15(4), 870–878. https://doi.org/10.1006/nimg.2001.1037

German, D. C., Walker, B. S., Manaye, K., Smith, W. K., Woodward, D. J., & North, A. J. (1988). The human locus coeruleus: Computer reconstruction of cellular distribution. Journal of Neuroscience, 8(5), 1776–1788. https://doi.org/10.1523/JNEUROSCI.08-05-01776.1988

Gilzenrat, M. S., Nieuwenhuis, S., Jepma, M., & Cohen, J. D. (2010). Pupil diameter tracks changes in control state predicted by the adaptive gain theory of locus coeruleus function. Cognitive, Affective & Behavioral Neuroscience, 10(2), 252–269. https://doi.org/10.3758/CABN.10.2.252

Glennon, E., Carcea, I., Martins, A. R. O., Multani, J., Shehu, I., Svirsky, M. A., & Froemke, R. C. (2019). Locus coeruleus activation accelerates perceptual learning. Brain Research, 1709, 39–49. https://doi.org/10.1016/j.brainres.2018.05.048

Glover, G. H., Li, T.-Q., & Ress, D. (2000). Image-based method for retrospective correction of physiological motion effects in fMRI: RETROICOR. Magnetic Resonance in Medicine, 44(1), 162–167. https://doi.org/10.1002/1522-2594(200007)44:1<162::AID-MRM23>3.0.CO;2-E

Greene, C. M., Bellgrove, M. A., Gill, M., & Robertson, I. H. (2009). Noradrenergic genotype predicts lapses in sustained attention. Neuropsychologia, 47(2), 591–594. https://doi.org/10.1016/j.neuropsychologia.2008.10.003

Grella, S. L., Neil, J. M., Edison, H. T., Strong, V. D., Odintsova, I. V., Walling, S. G., … Harley, C. W. (2019). Locus Coeruleus Phasic, But Not Tonic, Activation Initiates Global Remapping in a Familiar Environment. Journal of Neuroscience, 39(3), 445–455. https://doi.org/10.1523/JNEUROSCI.1956-18.2018

Halliday, G. M., Fedorow, H., Rickert, C. H., Gerlach, M., Riederer, P., & Double, K. L. (2006). Evidence for specific phases in the development of human neuromelanin. Journal of Neural Transmission, 113(6), 721–728. https://doi.org/10.1007/s00702-006-0449-y

Hausser, J., Strimmer, K., & Strimmer, M. K. (2012). Package ‘entropy’ for R.

Jepma, M., & Nieuwenhuis, S. (2010). Pupil Diameter Predicts Changes in the Exploration–Exploitation Trade-off: Evidence for the Adaptive Gain Theory. Journal of Cognitive Neuroscience, 23(7), 1587–1596. https://doi.org/10.1162/jocn.2010.21548

Jo, H. J., Saad, Z. S., Simmons, W. K., Milbury, L. A., & Cox, R. W. (2010). Mapping Sources of Correlation in Resting State FMRI, with Artifact Detection and Removal. NeuroImage, 52(2), 571–582. https://doi.org/10.1016/j.neuroimage.2010.04.246

Jones, B. E., & Moore, R. Y. (1977). Ascending projections of the locus coeruleus in the rat. II. Autoradiographic study. Brain Research, 127(1), 23–53. https://doi.org/10.1016/0006-8993(77)90378-X

Jones, B. E., & Yang, T.-Z. (1985). The efferent projections from the reticular formation and the locus coeruleus studied by anterograde and retrograde axonal transport in the rat. Journal of Comparative Neurology, 242(1), 56–92. https://doi.org/10.1002/cne.902420105

Joshi, S., Li, Y., Kalwani, R., & Gold, J. I. (2016). Relationships between pupil diameter and neuronal activity in the locus coeruleus, colliculi, and cingulate cortex. Neuron, 89(1), 221–234. https://doi.org/10.1016/j.neuron.2015.11.028

Keren, N. I., Lozar, C. T., Harris, K. C., Morgan, P. S., & Eckert, M. A. (2009). In vivo mapping of the human locus coeruleus. NeuroImage, 47(4), 1261–1267. https://doi.org/10.1016/j.neuroimage.2009.06.012

Keren, N. I., Taheri, S., Vazey, E. M., Morgan, P. S., Granholm, A.-C. E., Aston-Jones, G. S., & Eckert, M. A. (2015). Histologic validation of locus coeruleus MRI contrast in post-mortem tissue. NeuroImage, 113, 235–245. https://doi.org/10.1016/j.neuroimage.2015.03.020

Kline, R. L., Zhang, S., Farr, O. M., Hu, S., Zaborszky, L., Samanez-Larkin, G. R., & Li, C.-S. R. (2016). The Effects of Methylphenidate on Resting-State Functional Connectivity of the Basal Nucleus of Meynert, Locus Coeruleus, and Ventral Tegmental Area in Healthy Adults. Frontiers in Human Neuroscience, 10. https://doi.org/10.3389/fnhum.2016.00149

Köhler, S., Wagner, G., & Bär, K.-J. (2019). Activation of brainstem and midbrain nuclei during cognitive control in medicated patients with schizophrenia. Human Brain Mapping, 40(1), 202–213. https://doi.org/10.1002/hbm.24365

Krebs, R. M., Park, H. R., Bombeke, K., & Boehler, C. N. (2018). Modulation of locus coeruleus activity by novel oddball stimuli. Brain Imaging and Behavior, 12(2), 577–584. http://doi.org/10.1007/s11682-017-9700-4

Kundu, P., Brenowitz, N. D., Voon, V., Worbe, Y., Vértes, P. E., Inati, S. J., … Bullmore, E. T. (2013). Integrated strategy for improving functional connectivity mapping using multiecho fMRI. Proceedings of the National Academy of Sciences, 110(40), 16187–16192. https://doi.org/10.1073/pnas.1301725110

Kundu, P., Inati, S. J., Evans, J. W., Luh, W.-M., & Bandettini, P. A. (2012). Differentiating BOLD and Non-BOLD Signals in fMRI Time Series Using Multi-Echo EPI. Neuroimage, 60(3), 1759–1770. https://doi.org/10.1016/j.neuroimage.2011.12.028

Kundu, P., Voon, V., Balchandani, P., Lombardo, M. V., Poser, B. A., & Bandettini, P. A. (2017). Multi-echo fMRI: A review of applications in fMRI denoising and analysis of BOLD signals. NeuroImage, 154, 59–80. https://doi.org/10.1016/j.neuroimage.2017.03.033

Ladd, M. E., Bachert, P., Meyerspeer, M., Moser, E., Nagel, A. M., Norris, D. G., … Zaiss, M. (2018). Pros and cons of ultra-high-field MRI/MRS for human application. Progress in Nuclear Magnetic Resonance Spectroscopy, 109, 1–50. https://doi.org/10.1016/j.pnmrs.2018.06.001

Liu, K. Y., Acosta-Cabronero, J., Cardenas-Blanco, A., Loane, C., Berry, A. J., Betts, M. J., … Hämmerer, D. (2019). In vivo visualization of age-related differences in the locus coeruleus. Neurobiology of Aging, 74, 101–111. https://doi.org/10.1016/j.neurobiolaging.2018.10.014

Liu, K. Y., Marijatta, F., Hämmerer, D., Acosta-Cabronero, J., Düzel, E., & Howard, R. J. (2017). Magnetic resonance imaging of the human locus coeruleus: A systematic review. Neuroscience & Biobehavioral Reviews, 83, 325–355. https://doi.org/10.1016/j.neubiorev.2017.10.023

Liu, T. T., Nalci, A., & Falahpour, M. (2017). The Global Signal in fMRI: Nuisance or Information? NeuroImage, 150, 213–229. https://doi.org/10.1016/j.neuroimage.2017.02.036

Ljubojevic, V., Luu, P., & Rosa, E. D. (2014). Cholinergic Contributions to Supramodal Attentional Processes in Rats. Journal of Neuroscience, 34(6), 2264–2275. https://doi.org/10.1523/JNEUROSCI.1024-13.2014

Lombardo, M. V., Auyeung, B., Holt, R. J., Waldman, J., Ruigrok, A. N. V., Mooney, N., … Kundu, P. (2016). Improving effect size estimation and statistical power with multi-echo fMRI and its impact on understanding the neural systems supporting mentalizing. Neuroimage, 142, 55–66. https://doi.org/10.1016/j.neuroimage.2016.07.022

Loughlin, S. E., Foote, S. L., & Bloom, F. E. (1986). Efferent projections of nucleus locus coeruleus: Topographic organization of cells of origin demonstrated by three-dimensional reconstruction. Neuroscience, 18(2), 291–306. https://doi.org/10.1016/0306-4522(86)90155-7

Loughlin, Sandra E., Foote, S. L., & Fallon, J. H. (1982). Locus coeruleus projections to cortex: Topography, morphology and collateralization. Brain Research Bulletin, 9(1), 287–294. https://doi.org/10.1016/0361-9230(82)90142-3

Luppi, P.-H., Aston-Jones, G., Akaoka, H., Chouvet, G., & Jouvet, M. (1995). Afferent projections to the rat locus coeruleus demonstrated by retrograde and anterograde tracing with cholera-toxin B subunit and Phaseolus vulgaris leucoagglutinin. Neuroscience, 65(1), 119–160. https://doi.org/10.1016/0306-4522(94)00481-J

Mann, D. M., Yates, P. O., & Hawkes, J. (1983). The pathology of the human locus ceruleus. Clinical Neuropathology, 2(1), 1–7.

Marien, M. R., Colpaert, F. C., & Rosenquist, A. C. (2004). Noradrenergic mechanisms in neurodegenerative diseases: A theory. Brain Research Reviews, 45(1), 38–78. https://doi.org/10.1016/j.brainresrev.2004.02.002

Markello, R. D., Spreng, R. N., Luh, W.-M., Anderson, A. K., & De Rosa, E. (2018). Segregation of the human basal forebrain using resting state functional MRI. NeuroImage, 173, 287–297. https://doi.org/10.1016/j.neuroimage.2018.02.042

Mather, M., Clewett, D., Sakaki, M., & Harley, C. W. (2016). Norepinephrine ignites local hot spots of neuronal excitation: How arousal amplifies selectivity in perception and memory. The Behavioral and Brain Sciences, 39, e200. https://doi.org/10.1017/S0140525X15000667

Mather, M., & Harley, C. W. (2016). The Locus Coeruleus: Essential for Maintaining Cognitive Function and the Aging Brain. Trends in Cognitive Sciences, 20(3), 214–226. https://doi.org/10.1016/j.tics.2016.01.001

Mesulam, M.-M. (2013). Cholinergic Circuitry of the Human Nucleus Basalis and Its Fate in Alzheimer’s Disease. The Journal of Comparative Neurology, 521(18), 4124–4144. https://doi.org/10.1002/cne.23415

Mikl, M., Mareček, R., Hluštík, P., Pavlicová, M., Drastich, A., Chlebus, P., … Krupa, P. (2008). Effects of spatial smoothing on fMRI group inferences. Magnetic Resonance Imaging, 26(4), 490–503. https://doi.org/10.1016/j.mri.2007.08.006

Mohanty, A., Gitelman, D. R., Small, D. M., & Mesulam, M. M. (2008). The Spatial Attention Network Interacts with Limbic and Monoaminergic Systems to Modulate Motivation-Induced Attention Shifts. Cerebral Cortex (New York, NY), 18(11), 2604–2613. https://doi.org/10.1093/cercor/bhn021

Mouton, P. R., Pakkenberg, B., Gundersen, H. J. G., & Price, D. L. (1994). Absolute number and size of pigmented locus coeruleus neurons in young and aged individuals. Journal of Chemical Neuroanatomy, 7(3), 185–190. https://doi.org/10.1016/0891-0618(94)90028-0

Murphy, P. R., O’Connell, R. G., O’Sullivan, M., Robertson, I. H., & Balsters, J. H. (2014). Pupil diameter covaries with BOLD activity in human locus coeruleus. Human Brain Mapping, 35(8), 4140–4154. https://doi.org/10.1002/hbm.22466

Neary, D. (2008). Chapter 19 - Brain Stem. In S. Standring & A. R. Crossman (Eds.), Gray’s Anatomy, (Ed. 40, pp. 275–296). Edinburgh: Churchill Livingstone.

Nieuwenhuis, S., van Nieuwpoort, I. C., Veltman, D. J., & Drent, M. L. (2007). Effects of the noradrenergic agonist clonidine on temporal and spatial attention. Psychopharmacology, 193(2), 261–269. https://doi.org/10.1007/s00213-007-0770-7

Park, H. W., Ro, Y. M., & Cho, Z. H. (1988). Measurement of the magnetic susceptibility effect in high-field NMR imaging. Physics in Medicine and Biology, 33(3), 339–349. https://doi.org/10.1088/0031-9155/33/3/003

Power, J. D., Barnes, K. A., Snyder, A. Z., Schlaggar, B. L., & Petersen, S. E. (2012). Spurious but systematic correlations in functional connectivity MRI networks arise from subject motion. Neuroimage, 59(3), 2142–2154. https://doi.org/10.1016/j.neuroimage.2011.10.018

Power, J. D., Plitt, M., Gotts, S. J., Kundu, P., Voon, V., Bandettini, P. A., & Martin, A. (2018). Ridding fMRI data of motion-related influences: Removal of signals with distinct spatial and physical bases in multiecho data. Proceedings of the National Academy of Sciences, 115(9), E2105–E2114. https://doi.org/10.1073/pnas.1720985115

Preibisch, C., Castrillón G., J. G., Bührer, M., & Riedl, V. (2015). Evaluation of Multiband EPI Acquisitions for Resting State fMRI. PLoS ONE, 10(9). https://doi.org/10.1371/journal.pone.0136961

Pruim, R. H. R., Mennes, M., van Rooij, D., Llera, A., Buitelaar, J. K., & Beckmann, C. F. (2015). ICA-AROMA: A robust ICA-based strategy for removing motion artifacts from fMRI data. NeuroImage, 112, 267–277. https://doi.org/10.1016/j.neuroimage.2015.02.064

Ryan, P. J., Ma, S., Olucha-Bordonau, F. E., & Gundlach, A. L. (2011). Nucleus incertus—An emerging modulatory role in arousal, stress and memory. Neuroscience & Biobehavioral Reviews, 35(6), 1326–1341. https://doi.org/10.1016/j.neubiorev.2011.02.004

Saad, Z. S., Glen, D. R., Chen, G., Beauchamp, M. S., Desai, R., & Cox, R. W. (2009). A New Method for Improving Functional-to-Structural MRI Alignment using Local Pearson Correlation. NeuroImage, 44(3), 839–848. https://doi.org/10.1016/j.neuroimage.2008.09.037

Samuels, E. R., & Szabadi, E. (2008). Functional Neuroanatomy of the Noradrenergic Locus Coeruleus: Its Roles in the Regulation of Arousal and Autonomic Function Part I: Principles of Functional Organisation. Current Neuropharmacology, 6(3), 235–253. http://doi.org/10.2174/157015908785777229

Sara. S. J. (2009). The locus coeruleus and noradrenergic modulation of cognition. Nature Reviews Neuroscience, 10(3), 211–223.

Sarter, M., Hasselmo, M. E., Bruno, J. P., & Givens, B. (2005). Unraveling the attentional functions of cortical cholinergic inputs: Interactions between signal-driven and cognitive modulation of signal detection. Brain Research Reviews, 48(1), 98–111. https://doi.org/10.1016/j.brainresrev.2004.08.006

Sasaki, M., Shibata, E., Tohyama, K., Takahashi, J., Otsuka, K., Tsuchiya, K., … Sakai, A. (2006). Neuromelanin magnetic resonance imaging of locus ceruleus and substantia nigra in Parkinson’s disease. NeuroReport, 17(11), 1215. https://doi.org/10.1097/01.wnr.0000227984.84927.a7

Schwarz, L. A., & Luo, L. (2015). Organization of the Locus Coeruleus-Norepinephrine System. Current Biology, 25(21), R1051–R1056. https://doi.org/10.1016/j.cub.2015.09.039

Shine, J. M., Bissett, P. G., Bell, P. T., Koyejo, O., Balsters, J. H., Gorgolewski, K. J., … Poldrack, R. A. (2016). The Dynamics of Functional Brain Networks: Integrated Network States during Cognitive Task Performance. Neuron, 92(2), 544–554. https://doi.org/10.1016/j.neuron.2016.09.018

Song, A. H., Kucyi, A., Napadow, V., Brown, E. N., Loggia, M. L., & Akeju, O. (2017). Pharmacological Modulation of Noradrenergic Arousal Circuitry Disrupts Functional Connectivity of the Locus Ceruleus in Humans. The Journal of Neuroscience, 37(29), 6938–6945. https://doi.org/10.1523/JNEUROSCI.0446-17.2017

Sterpenich, V., D’Argembeau, A., Desseilles, M., Balteau, E., Albouy, G., Vandewalle, G., … Maquet, P. (2006). The Locus Ceruleus Is Involved in the Successful Retrieval of Emotional Memories in Humans. Journal of Neuroscience, 26(28), 7416–7423. https://doi.org/10.1523/JNEUROSCI.1001-06.2006

Sulzer, D., Cassidy, C., Horga, G., Kang, U. J., Fahn, S., Casella, L., … Zecca, L. (2018). Neuromelanin detection by magnetic resonance imaging (MRI) and its promise as a biomarker for Parkinson’s disease. NPJ Parkinson’s Disease, 4(1), 1–13. https://doi.org/10.1038/s41531-018-0047-3

Swallow, K. M., Braver, T. S., Snyder, A. Z., Speer, N. K., & Zacks, J. M. (2003). Reliability of functional localization using fMRI. NeuroImage, 20(3), 1561–1577.

Swallow, K. M., Jiang, Y. V., & Riley, E. B. (2019). Target detection increases pupil diameter and enhances memory for background scenes during multi-tasking. Scientific Reports, 9(1), 1–13. https://doi.org/10.1038/s41598-019-41658-4

Szőnyi, A., Sos, K. E., Nyilas, R., Schlingloff, D., Domonkos, A., Takács, V. T., … Nyiri, G. (2019). Brainstem nucleus incertus controls contextual memory formation. Science, 364(6442), eaaw0445. https://doi.org/10.1126/science.aaw0445

Tona, K.-D., Keuken, M. C., de Rover, M., Lakke, E., Forstmann, B. U., Nieuwenhuis, S., & van Osch, M. J. P. (2017). In vivo visualization of the locus coeruleus in humans: Quantifying the test–retest reliability. Brain Structure and Function, 222(9), 4203–4217. https://doi.org/10.1007/s00429-017-1464-5

Uematsu, A., Tan, B. Z., & Johansen, J. P. (2015). Projection specificity in heterogeneous locus coeruleus cell populations: Implications for learning and memory. Learning & Memory, 22(9), 444–451. https://doi.org/10.1101/lm.037283.114

Uğurbil, K., Garwood, M., Ellermann, J., Hendrich, K., Hinke, R., Hu, X., … Ogawa, S. (1993). Imaging at high magnetic fields: Initial experiences at 4 T. Magnetic Resonance Quarterly, 9(4), 259–277.

van den Heuvel, M. P., & Hulshoff Pol, H. E. (2010). Exploring the brain network: A review on resting-state fMRI functional connectivity. European Neuropsychopharmacology: The Journal of the European College of Neuropsychopharmacology, 20(8), 519–534. https://doi.org/10.1016/j.euroneuro.2010.03.008

Vijayashankar, N., & Brody, H. (1979). A Quantitative Study of the Pigmented Neurons in the Nuclei Locus Coeruleus and Subcoeruleus in Man as Related to Aging. Journal of Neuropathology & Experimental Neurology, 38(5), 490–497. https://doi.org/10.1097/00005072-197909000-00004

Wakamatsu, K., Tabuchi, K., Ojika, M., Zucca, F. A., Zecca, L., & Ito, S. (2015). Norepinephrine and its metabolites are involved in the synthesis of neuromelanin derived from the locus coeruleus. Journal of Neurochemistry, 135(4), 768–776. https://doi.org/10.1111/jnc.13237

Windischberger, C., Langenberger, H., Sycha, T., Tschernko, E. M., Fuchsjäger-Mayerl, G., Schmetterer, L., & Moser, E. (2002). On the origin of respiratory artifacts in BOLD-EPI of the human brain. Magnetic Resonance Imaging, 20(8), 575–582. https://doi.org/10.1016/S0730-725X(02)00563-5

Worsley, K. J., Marrett, S., Neelin, P., Vandal, A. C., Friston, K. J., & Evans, A. C. (1996). A unified statistical approach for determining significant signals in images of cerebral activation. Human Brain Mapping, 4(1), 58–73. https://doi.org/10.1002/(SICI)1097-0193(1996)4:1<58::AID-HBM4>3.0.CO;2-O

Yarkoni, T. (2009). Big Correlations in Little Studies: Inflated fMRI Correlations Reflect Low Statistical Power—Commentary on Vul et al. (2009). Perspectives on Psychological Science, 4(3), 294–298. https://doi.org/10.1111/j.1745-6924.2009.01127.x

Yu, A. J., & Dayan, P. (2005). Uncertainty, Neuromodulation, and Attention. Neuron, 46(4), 681–692. https://doi.org/10.1016/j.neuron.2005.04.026

Zhang, S., Hu, S., Chao, H. H., & Li, C.-S. R. (2016). Resting-State Functional Connectivity of the Locus Coeruleus in Humans: In Comparison with the Ventral Tegmental Area/Substantia Nigra Pars Compacta and the Effects of Age. Cerebral Cortex (New York, NY), 26(8), 3413–3427. https://doi.org/10.1093/cercor/bhv172

