## Supplementary Information for "Estimates of locus coeruleus function with functional magnetic resonance imaging are influenced by localization approaches and the use of multi-echo data"

RUNNING HEAD: ESTIMATES OF LC FUNCTION

+ To whom correspondence should be addressed.
Khena Swallow
Department of Psychology
Cornell University
211 Uris Hall
Ithaca, NY 14853

Declaration of Interests: None.

### Supplemental Materials

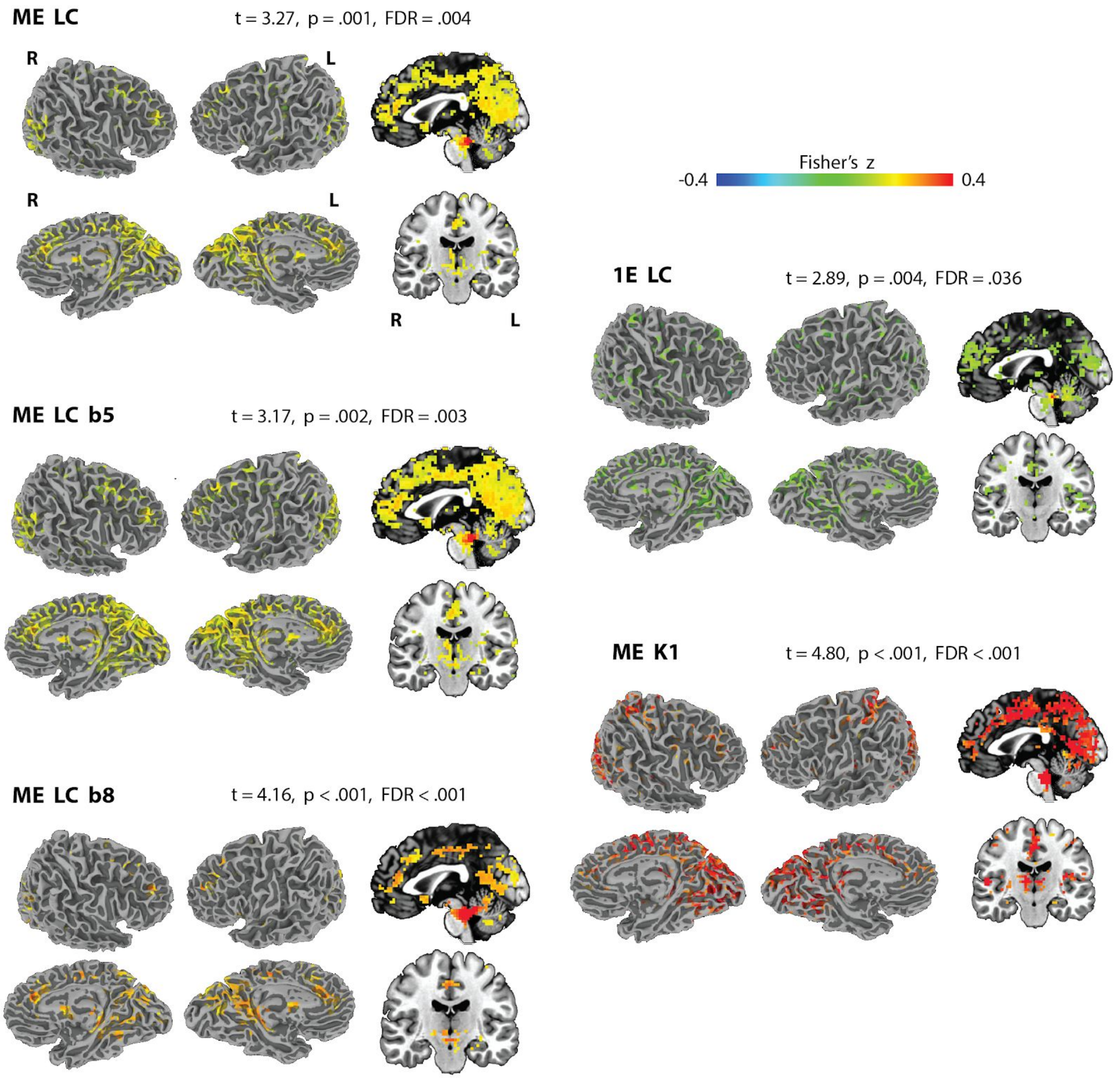


**Supplementary Figure S1.** Surface maps and sagittal and coronal slices of the brainstem illustrating the iFC maps generated by multiple pipelines and ROIs, including blurring to FWHM of 8 mm (ME LC b8), and that have been thresholded to the maximal Euler characteristic.

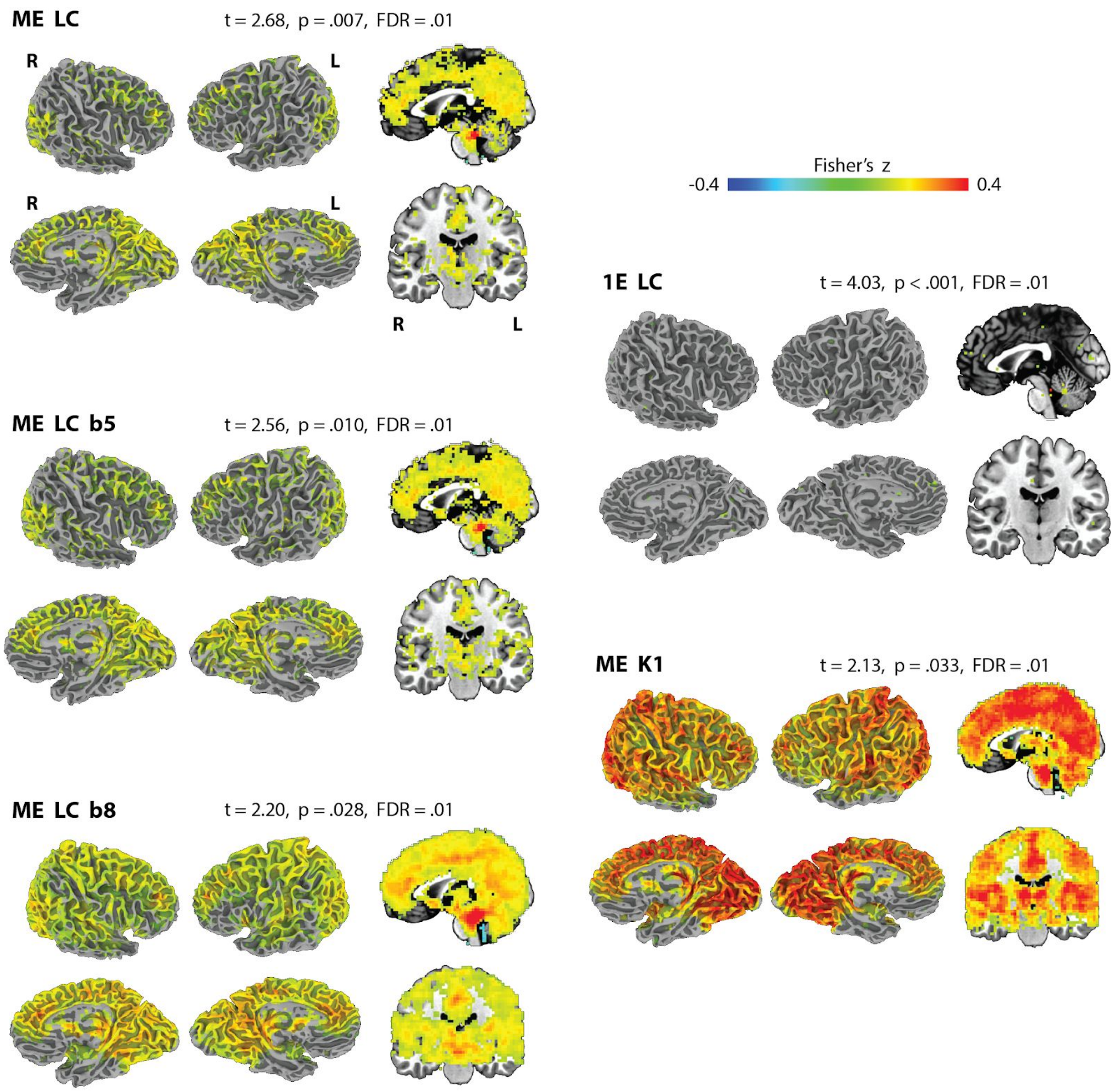


**Supplementary Figure S2.** Surface maps and sagittal and coronal slices of the brainstem illustrating the iFC maps generated by multiple pipelines and ROIs, including blurring the seed to FWHM of 8, and that have been thresholded to an FDR of .01.

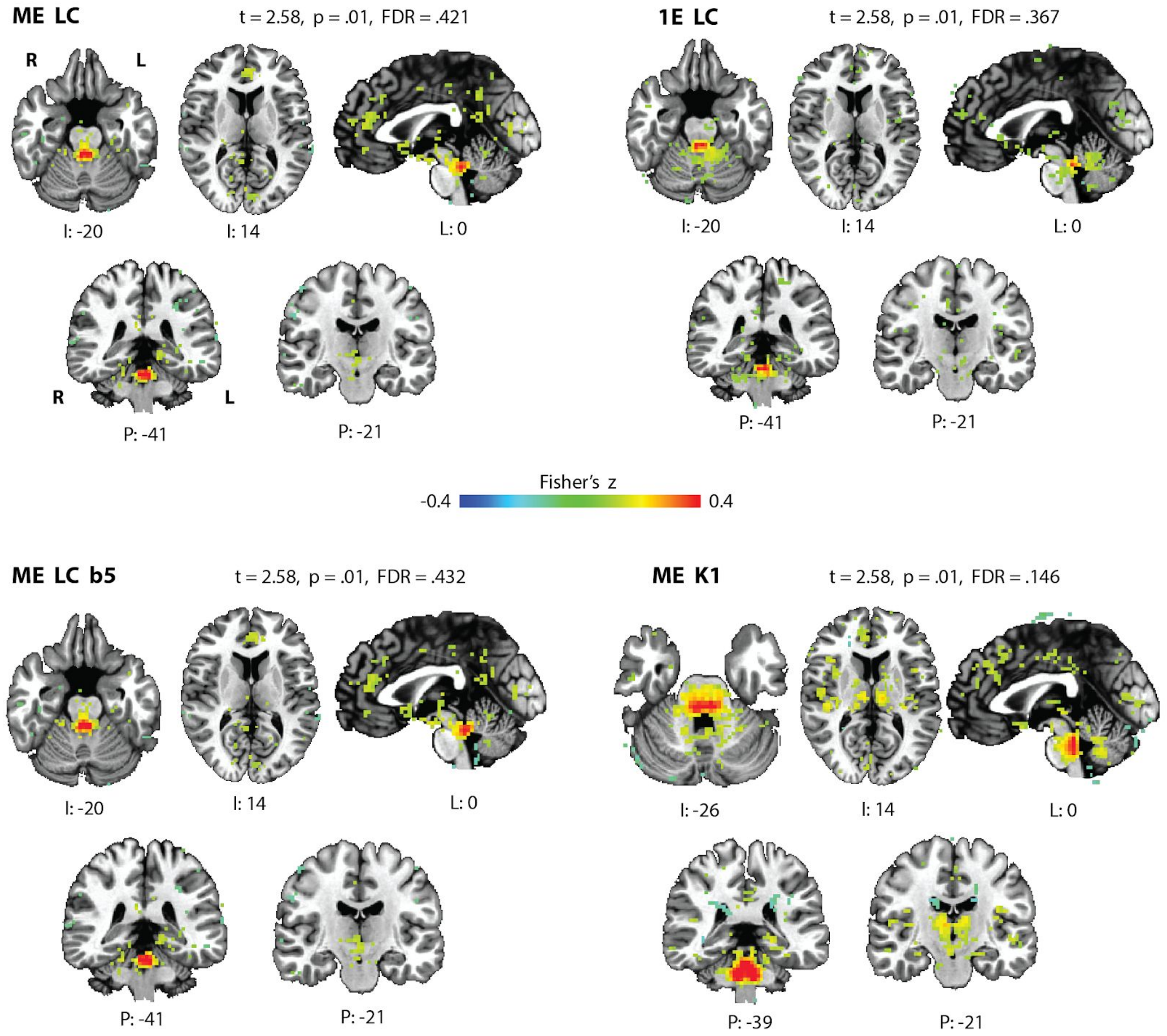


**Supplementary Figure S3.** iFC maps when the four main pipelines include global signal regression instead of white matter regression. Maps have been thresholded at p = .01; most voxels did not survive thresholding at an FDR of .05.

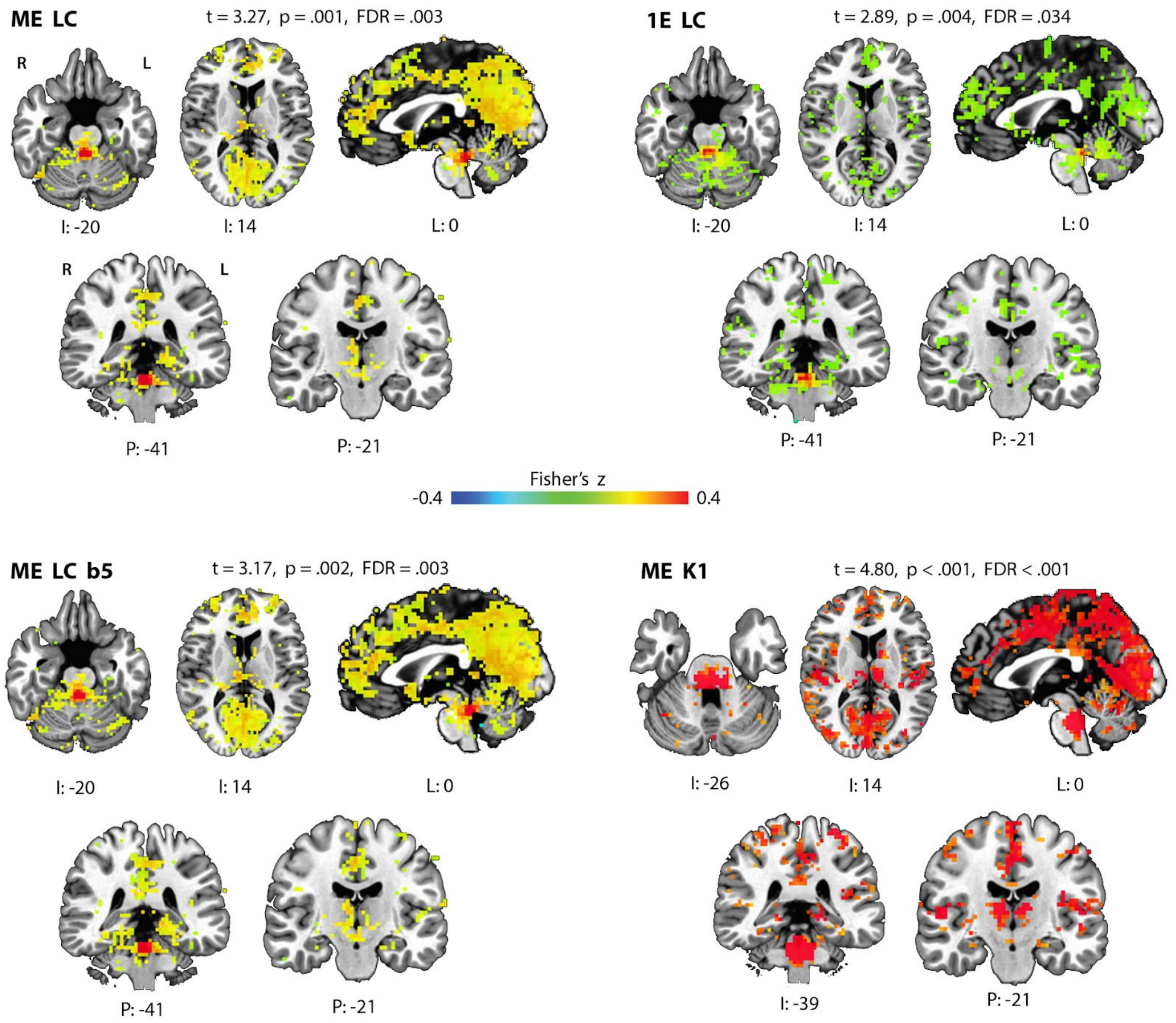


**Supplementary Figure S4.** iFC maps when the four main pipelines do not include white matter regression. Maps have been thresholded at the same t-statistic as in Figure 4.

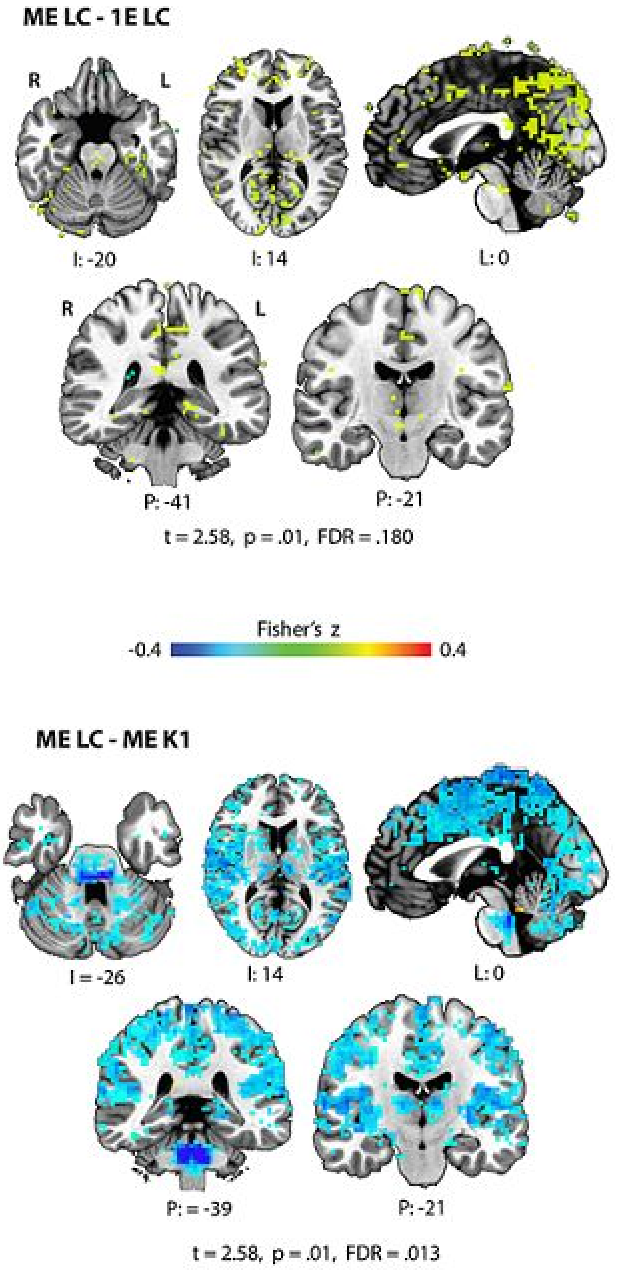


**Supplementary Figure S5.** Two-sample t-tests contrasting the ME LC pipeline against the other pipelines. Maps were thresholded at p = .01, at which point none of the voxels in the contrast against ME LC b5 survived. Note the pontine area (sagittal slice) that is greater for ME LC than for ME K1 that results from a linear transformation of the functional data, where the Keren atlas was non-linearly warped to the brainstem and thereby results in different voxels being included in the locus coeruleus ROI.
